# Human-specific morphoregulatory signatures in basal radial glia characterize neocortex evolution

**DOI:** 10.1101/2025.03.12.642828

**Authors:** Theresa M. Schütze, Nora Ditzer, Silvia Vangelisti, Annika Kolodziejczyk, Emanuele Capra, Ilaria Chiaradia, Jula Peters, Maximilian Krause, Christina Eugster, Razvan P. Derihaci, Cahit Birdir, Ulrich Martin, Pauline Wimberger, Katherine R. Long, Madeline Lancaster, Nereo Kalebic, Boyan Bonev, Mareike Albert

## Abstract

As the seat of our cognition, the human neocortex is an object of immense fascination. Human neocortex expansion during evolution has been attributed to an increase in the proliferative capacity of neural progenitor cells during development, particularly basal radial glia (bRG). Despite their evolutionary relevance, the genomic changes driving human-specific bRG biology remain uncharacterized. We used comparative chromatin and transcriptional profiling of neural progenitor cells isolated from gorilla, chimpanzee and human cerebral organoids to identify cis-regulatory elements that have gained activity in humans. Focusing specifically on bRG, we discovered that morphoregulatory enhancer activity and gene expression signatures distinguish human bRG from other great apes. Functional analysis of the morphoregulatory genes *FAM107A* and *CNGA3* in human organoids revealed that these genes are required for the morphological complexity of human bRG. Taken together, our interspecies comparison of basal radial glia suggests that human-specific morphoregulatory signatures characterize neocortex evolution.

## Main Text

As the seat of higher-level cognitive functions, including speech, self-reflection, and long-term planning (*1, 2*), the neocortex is central to the experience of being human. In the course of primate evolution, the neocortex has undergone a number of significant changes (*3, 4*), including a drastic increase in size and neuron number, both of which peak in humans (*5*). Neocortical neurons arise embryonically from three main classes of neural progenitor cells: (i) apical radial glia (aRG) that divide in the ventricular zone of the developing cortex, (ii) basal intermediate progenitors (bIPs) without ventricular contact that divide in the subventricular zone (SVZ), and (iii) basal (or outer) radial glia (bRG/oRG) that also divide in the SVZ and have repeatedly been implicated in the evolution of neocortex size. The abundance (*6-9*) and proliferative potential (*10-12*) of bRG are tightly linked to increased neuron numbers in the adult neocortex and overall neocortex size. Key factors underlying bRG function include their morphological complexity (*11, 13*), lineage relationships (*11, 14, 15*) and altered cell metabolism (*16*), all of which correlate with larger neocortex size across species.

The human-specific genomic changes underlying human bRG biology remain to be elucidated, promising unique insights into the mechanisms that ultimately provide the basis for higher cognitive functions. While a limited number of human-specific genes affecting bRG proliferative potential and neocortex size have been described (*3, 4, 17*), a large part of great ape species-specific biology likely originates in the differential regulation of shared genes orchestrated by cis-regulatory elements (CREs) (*18-20*). Inter-species differences in CRE activity may alter the expression of associated target genes, affecting radial glia biology, as shown for the Wnt receptor FZD8 (*21*) and the growth factor EPIREGULIN (*22*). Epigenomic comparisons of fetal brain tissue from different species have identified candidate CREs, termed human gain enhancers (HGEs), with a potential role in neocortex evolution (*23, 24*). Yet, since great ape fetal tissue is not accessible, truly human-specific changes of bRG CRE activity have not been elucidated. Recent advances in induced pluripotent stem cells (iPSCs) from great apes has allowed epigenetic comparisons of cultured neural progenitor cells and neurons (*25*), which, however, provide a limited recapitulation of *in vivo* development due to the inability to recapitulate tissue morphology in 2D cultures. Great ape cerebral organoid models can offer additional insights into human-specific enhancer activity (*26*), in particular, as cerebral organoids contain bRG (*27, 28*). Using 3D organoid models, here, we set out to identify CRE activity patterns specific to human bRG that are likely to contribute to their extensive proliferative potential, neuronal output, and, ultimately, human neocortex expansion.

### Cell type-specific ATAC-seq reveals distinct molecular profiles of great ape bRG

To characterize human-specific gene-regulatory signatures in bRG, we studied chromatin accessibility and gene expression in aRG, bRG, bIPs, and neurons during great ape cortical development (Fig. 1A). We generated cerebral organoids (CeOs) (*29*) from gorilla, chimpanzee, and human iPSCs (Fig. 1B, C) (*30-32*), expressing the forebrain marker FOXG1 (fig. S1A) and representing comparable stages of cortical development (fig. S1B). Application of the lipophilic dye DiI to the surface of CeOs reveals the presence of bRG with characteristic basal processes (Fig. 1D; fig S1C), including previously described morphological variety (*11, 13*), confirming that CeOs can be used to study bRG.

**Fig. 1.**
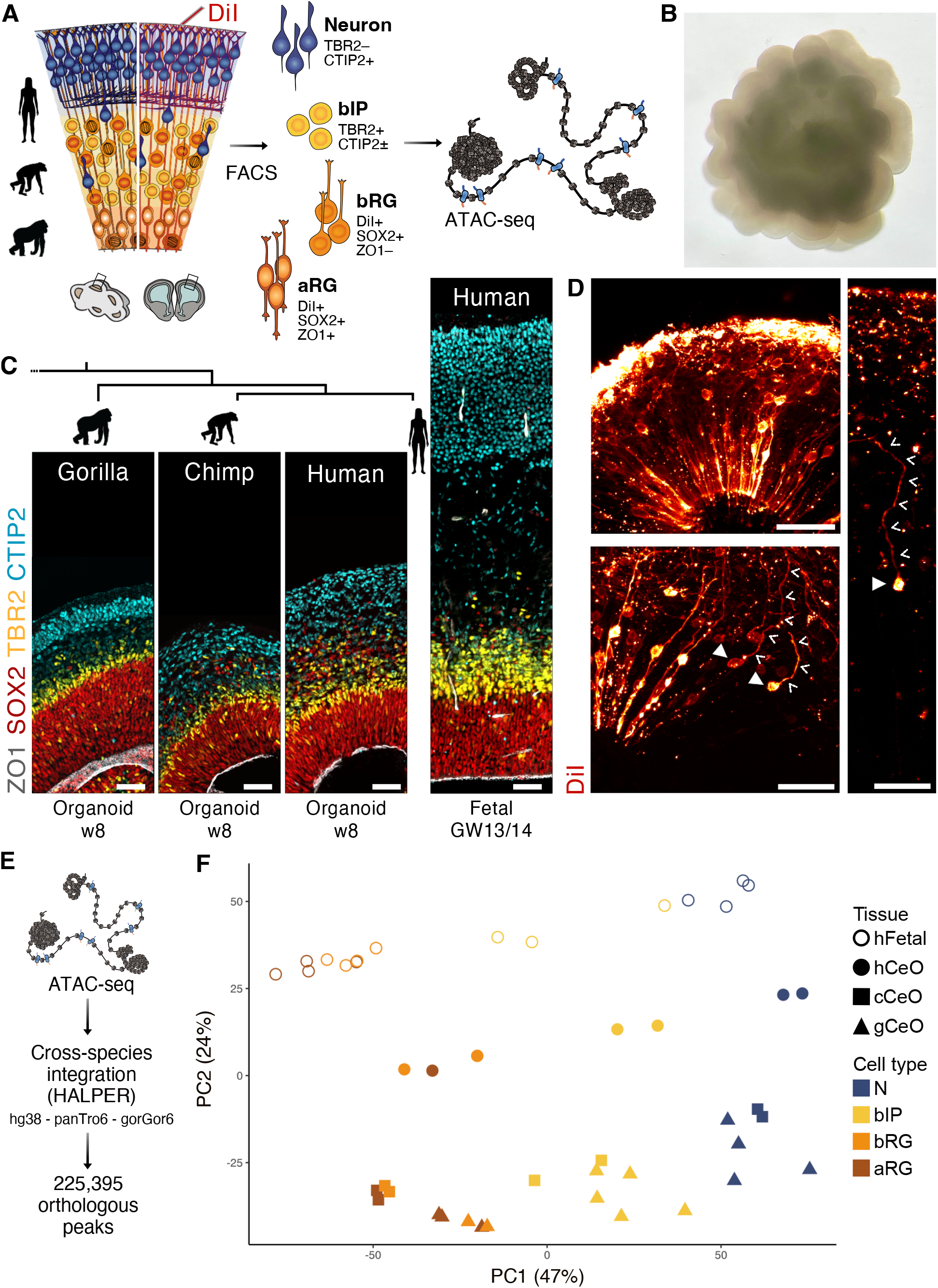
Cell type-specific ATAC-seq reveals distinct cellular profiles in great ape corticogenesis. (**A**) Scheme of the isolation of aRG, bRG, bIPs, and N from gorilla, chimpanzee, and human CeOs and human fetal cortex. The lipophilic dye DiI labels cells with processes reaching the basal side of the tissue, followed by tissue dissociation, immunocytochemistry and FACS. ATAC-seq is performed on all cell populations. Each data point with the same symbol represents an independent experiment. (**B**) Brightfield image of a whole gorilla CeO at w8. (**C**) Immunohistochemistry for ZO1, SOX2, TBR2, and CTIP2 in gorilla, chimpanzee, and human CeOs and human fetal cortex. (**D**) Sparse DiI labeling reveals radial tissue morphology and highlights the presence of bRG cell bodies (filled arrowheads) and processes (open arrowheads). (**E**) ATAC-seq libraries are sequenced by next-generation sequencing, and orthologs for regions mapped to hg38, panTro6, and gorGor6 are determined using HALPER (*37*). (**F**) Principal component analysis of the 1,000 most variable regions shows a separation of cell types on PC1 (47%) and species on PC2 (24%). gCeO, cCeO, hCeO, gorilla, chimpanzee and human cerebral organoid; hFetal, human fetal cortex. Scale bars, 50 μm.

We isolated specific cell populations from CeOs at week 8 and human fetal cortex at gestation weeks 13/14 (Fig. 1A–C, fig. S1D, E), based on the presence/absence of ventricular contact, basal processes and marker protein expression (*10, 33, 34*). RT-qPCR was used to confirm the identity of sorted cell populations (fig. S1F–H). We assayed open chromatin using ATAC-seq (*35, 36*) and integrated all great ape data sets using HALPER (*37*), yielding 225,395 orthologous peaks (Fig. 1E). We observe expected patterns of cell type-specific accessibility at marker genes and clustering of cell types (fig. S2A, B). Principal component analysis reveals a separation of cell types on PC1, accounting for 47% of the variation in the data, with a trajectory from radial glia to bIPs to neurons, demonstrating that broad cellular signatures are conserved across hominids (Fig. 1F). In line with previous literature (*14, 38-41*), aRG and bRG are not distinctly separated at this developmental stage and molecular level (Fig. 1F; fig. S2B, C); rather bRG show an overall unique pattern of shared accessibility features with the other neural cell types (fig. S2C). PC2 accounts for 24% of the variation and appears to separate the species, revealing divergent species-specific molecular profiles of the different cell types, including human-specific features of bRG (Fig. 1F).

### Human bRG exhibit species-specific chromatin accessibility patterns linked to morphoregulatory genes

Next, we aimed to define the species-specific patterns of bRG chromatin accessibility indicated by multivariate analysis. We, therefore, identified differentially accessible distal chromatin peaks in human compared to chimpanzee and gorilla bRG isolated from CeOs (log2FC ≥ 1.5; p ≤ 0.05). To ensure *in vivo* developmental relevance, we further cross-referenced these peaks with regions accessible in bRG isolated from human fetal cortex (Fig. 2A). This analysis led to the identification of 599 candidate CREs with higher accessibility in human bRG, which we termed human bRG candidate CREs (hbRG cCREs) (Fig. 2A, B; Data S1). These hbRG cCREs fall into three broad clusters of shared accessibility with other cell types: cluster 1) shows ancestral accessibility in bIPs and neurons, gained in human aRG and bRG; cluster 2) shows strongly increased accessibility in human aRG and bRG, shared to a lesser degree with radial glia of both other great apes, while cluster 3) shares RG accessibility only with gorillas (Fig. 2C). Interestingly, clusters 2 and 3 also show some accessibility in human bIPs, indicating they may be more broadly relevant to human neural progenitor cell biology. Characterizing hbRG cCREs further, we found that many (63%) overlap with CRE signatures defined by the ENCODE Project (*42*) and three regions overlap with the VISTA enhancer database (*43, 44*) (fig. S2D). Human accelerated regions, repeatedly indicated to harbor significant gene-regulatory potential relevant to human brain evolution (*25, 45-47*), are represented among the hbRG cCREs (Fig. 2D) (genome-wide enrichment tested by Fisher’s exact test; p = 0.0509; odds ratio = 2.3), adding support to the notion that human accelerated regions are highly relevant in the context of human-specific neurodevelopmental gene regulation.

**Fig. 2.**
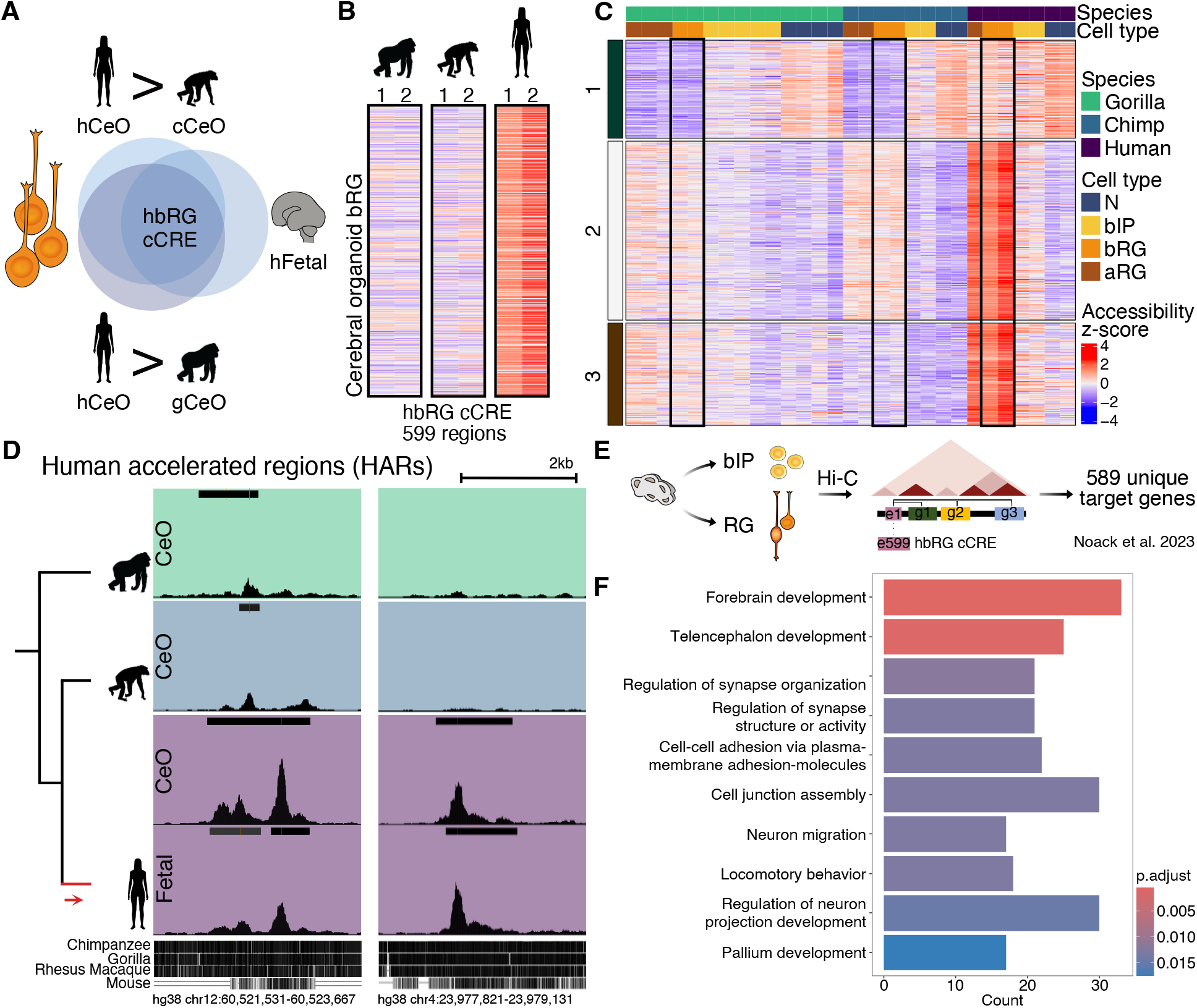
Human-specific patterns of chromatin accessibility in bRG link to cell morphology genes. (**A**) Human-increased chromatin accessibility in bRG candidate cis-regulatory elements (hbRG cCRE) is identified by overlapping regions found to be significantly more accessible (log2FC ≥ 1.5; p ≤ 0.05) in human bRG than in both chimpanzee and gorilla. Regions are benchmarked against peaks in human fetal bRG. (**B**) Accessibility z-scores of the 599 hbRG cCRE from (A) in the two replicates of gorilla, chimpanzee and human CeO bRG. (**C**) K-means clustering of the 599 hbRG cCRE reveals three broad patterns of accessibility across all CeO samples and cell types. Black boxes highlight gorilla, chimpanzee and human bRG. (**D**) Two examples of human accelerated regions that are found among the hbRG cCRE. Shown are merged bigWig tracks of at least two replicates, with standardized track height 0–3. Called peaks are represented by black bars in tracks. Cactus evolutionary sequence conservation and chromosome coordinates are indicated. (**E**) Candidate CREs were linked to target genes using cell type-specific Hi-C data from radial glia and bIPs (*48*), resulting in 589 unique target genes in physical contact with the 599 hbRG cCREs. (**F**) GO term analysis of target genes indicates enrichment of genes linked to forebrain development, regulation of cell-cell contacts and regulation of mature neural cell morphology. Top 10 terms displayed with p ≤ 0.05.

To explore the potential biological role of hbRG cCRE activity, we leveraged cell type-specific Hi-C data of cortical organoids (*48*) to link our candidate regions to their target genes. We found that the 599 regions target a total of 589 unique genes (Fig. 2E). GO term analysis revealed that after broader terms, such as ‘Forebrain development’, these genes appear to be predominantly linked to processes that regulate the function and morphology of mature neural cells, including ‘Regulation of neuron projection development’ (Fig. 2F). Human bRG may thus employ both ancestral neuronal CREs (Fig. 2C; cluster 1) as well as increased activity of ancestral radial glia CREs to express genes classically associated with controlling morphoregulatory processes in neurons.

### Human bRG-upregulated genes encode factors associated with cell projection and growth

To further investigate the possibility of altered morphoregulatory dynamics in human bRG, we turned to chimpanzee and human RNA-seq data (Fig. 3A). Principle component analysis (Fig. 3B) and gene expression correlation analysis (fig. S3A) mirror cell type- and species-specific patterns observed on chromatin level. Marker gene analysis confirms both regional and cell type identity at the transcriptomic level (fig. S3B, C). We found a total of 939 differentially expressed genes (log2FC ≥ 1; p ≤ 0.05) between chimpanzee and human bRG (Fig. 3C; Data S2, S3), whose expression pattern in hCeO is highly correlated to that of hFetal bRG (Fig. 3D). GO term analysis corroborates the pattern observed in hbRG cCRE target genes, with genes upregulated in human bRG predominantly enriched in processes regulating neural cell morphology (Fig. 3E). Of note, GO terms classifying processes such as the ‘Regulation of neuron or cell projections’ and ‘Axonogenesis’ are specific to genes upregulated in human bRG, and not detected for aRG and bIPs (fig. S3D, E). This suggests that altered morphoregulatory dynamics of cell projections are a human bRG-specific phenotype. Indeed, *PALMD*, previously identified as a crucial regulator of basal progenitor morphological complexity and proliferative potential in the context of neocortex evolution (*13*), is significantly upregulated in human bRG compared to chimpanzee bRG, but not in human aRG or bIPs (fig. S3F).

**Fig. 3.**
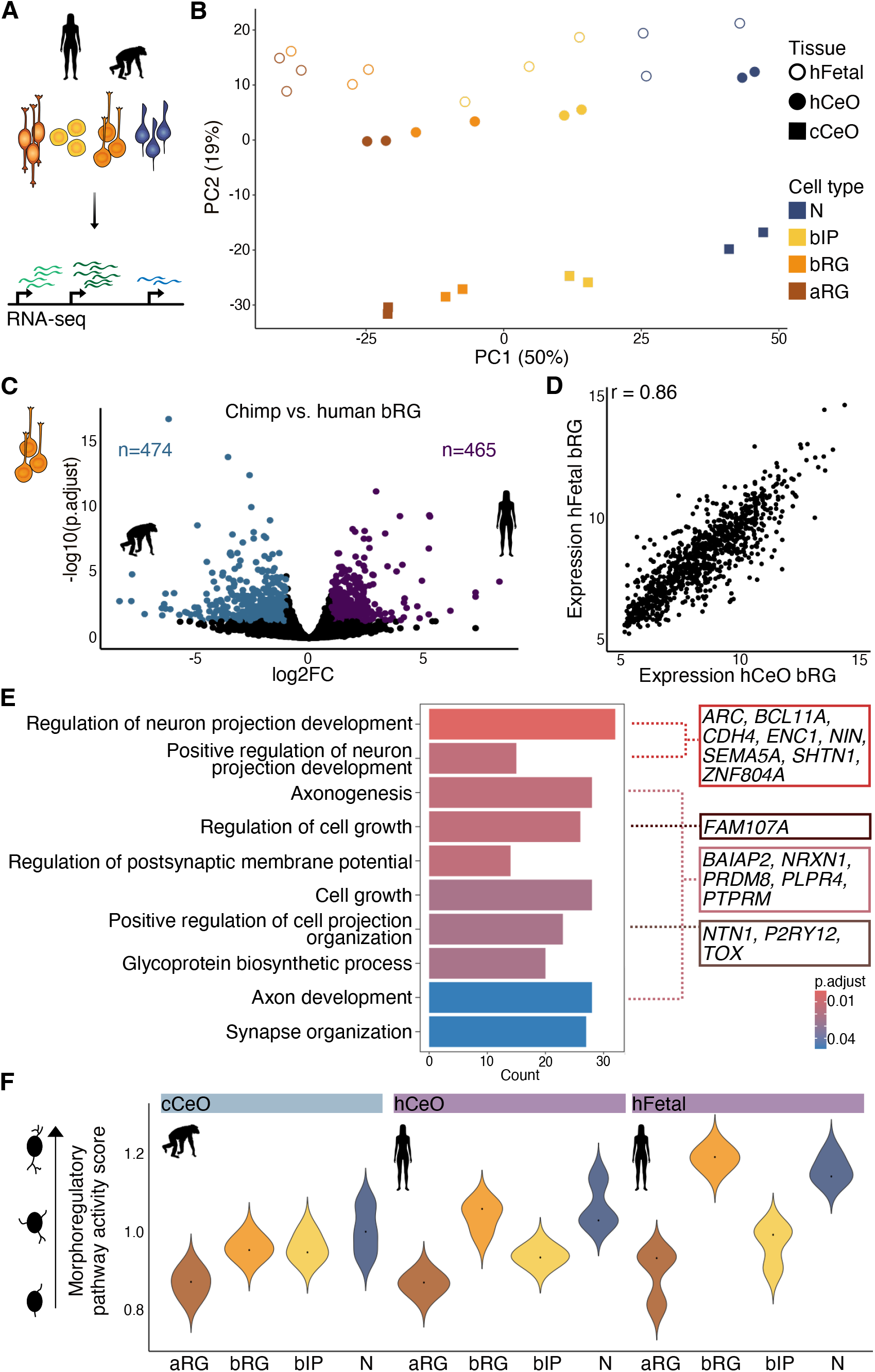
Genes more highly expressed in human than chimpanzee bRG indicate regulation of morphoregulatory pathways. (**A**) RNA-seq is performed on all sorted cell types from chimpanzee and human CeOs and human fetal cortex. (**B**) Principal Component analysis of 500 most variable genes shows a separation of cell types on PC1 (50%) and species on PC2 (19%). (**C**) Volcano plot showing genes significantly upregulated in chimpanzee (blue) and human (purple) bRG (FDR = 0.05; log2FC ≥ 1). (**D**) Correlation of variance stabilized expression of bRG DEGs in human CeO and fetal bRG. (**E**) GO term analysis of human bRG upregulated genes indicates enrichment of genes linked to the regulation of cell morphology. Top 10 terms displayed with p ≤ 0.05. Example genes are indicated on the right. (**F**) Morphoregulatory pathway activity score, adapted from (*50*) and calculated based on GO term gene groups, indicates higher morphoregulatory activity in human bRG compared to chimpanzee bRG.

Given that both hbRG cCRE target genes and human bRG upregulated genes indicate differential regulation of cell morphology, we explored this aspect in more detail. In analogy to previous approaches to determine a glycolysis score (*49*) or metabolic pathway activity (*50*), we calculated a ‘morphoregulatory pathway activity score’ based on the GO terms ‘Cell morphogenesis’, ‘Cell projection morphogenesis’ and ‘Cell projection organization’ (Fig. 3F; Materials and methods). In chimpanzees, this morphoregulatory pathway activity score increases from aRG to bRG and bIPs and then again to neurons, in line with the biological progression from a bipolar cell (aRG) to what in primates often are morphologically more complex basal progenitors (*11, 13*) to, finally, highly complex neurons. In humans, the pathway activity mirrors the general increase from aRG to basal progenitors to neurons. Of note, human bRG show a score almost identical to neurons, outstripping both human aRG and bIPs as well as chimpanzee neural progenitor cells. These results further underline the concept that altered morphoregulatory signatures are a key feature of human bRG biology and neocortex evolution.

### Human bRG show signatures of co-option from neurons on chromatin and transcriptome level

Finally, we sought to gain functional insight into hbRG cCREs and their morphoregulatory target genes. In analogy to the hbRG cCRE accessibility pattern (Fig. 2C), expression of the 589 target genes in humans falls into two broad clusters (fig. S4A). The first shows expression in the progenitor cell populations but foremost expression in neurons, including genes such as *STMN2* (fig. S4B), a microtubule-binding protein regulating neuronal neurite outgrowths (*51*), and *AUTS2* (fig. S4C), an autism risk gene previously shown to regulate actin dynamics and neurite formation in neurons (*52*). While *AUTS2* has recently been shown to increase basal progenitor production and neuron output in the mouse cortex (*53*), neither gene has so far been associated with regulating neural progenitor cell morphology–a highly interesting perspective given the evolutionary dynamics discovered here.

The second cluster of hbRG cCRE target genes shows high expression in neural progenitor cells, especially radial glia (fig. S4A). Interestingly, while the expression pattern of these genes is largely conserved in chimpanzees, a number of genes appear more highly expressed in chimpanzee neurons than neural progenitor cells (fig. S4D, chimpanzee cluster 1). Example genes include *PTPRM* (Fig. 3E; fig. S4E), a protein tyrosine phosphatase receptor shown to be critical for synapse formation (*54*), and *NRG1* (fig. S4F), a stimulator of neurite outgrowth in various neuron types (*55*). This suggested pattern of ancestral neuronal gene expression of a small number of genes in chimpanzees, in conjunction with the ancestral accessibility observed for some hbRG cCRE in chimpanzee and gorilla neurons (Fig. 2C, cluster 1) as well as the human-upregulated expression of genes associated with regulating neuronal cell morphology (Fig. 3E, F) leads us to propose that human bRG may have re-purposed ancestral neuronal CREs as well as neuronal genes to alter and potentially complexify their morphology.

### Increased regulatory potential of candidate human bRG morphoregulatory enhancers

As increased expression of morphoregulatory genes in basal progenitors was shown to be highly relevant for neocortex size evolution (*13*), we aimed to functionally investigate CRE-gene pairs falling into this category (fig. S4A, cluster 2). One such gene (Fig. 4A), *FAM107A*, an actin-binding protein suggested as a key component of the human bRG molecular signature (*40*), shows significantly higher expression in human compared to chimpanzee bRG (Fig. 4B; Data S2). The CRE targeting *FAM107A* (Fig. 4C) shows higher accessibility in human compared to gorilla and chimpanzee bRG. Luciferase reporter assays in N2A cells (Fig. 4D) revealed that only the human sequence of the *FAM107A* cCRE, and not the orthologous chimpanzee sequence, significantly drives reporter gene expression (Fig. 4E). This indicates that the human region has a higher gene regulatory potential than its chimpanzee ortholog.

**Fig. 4.**
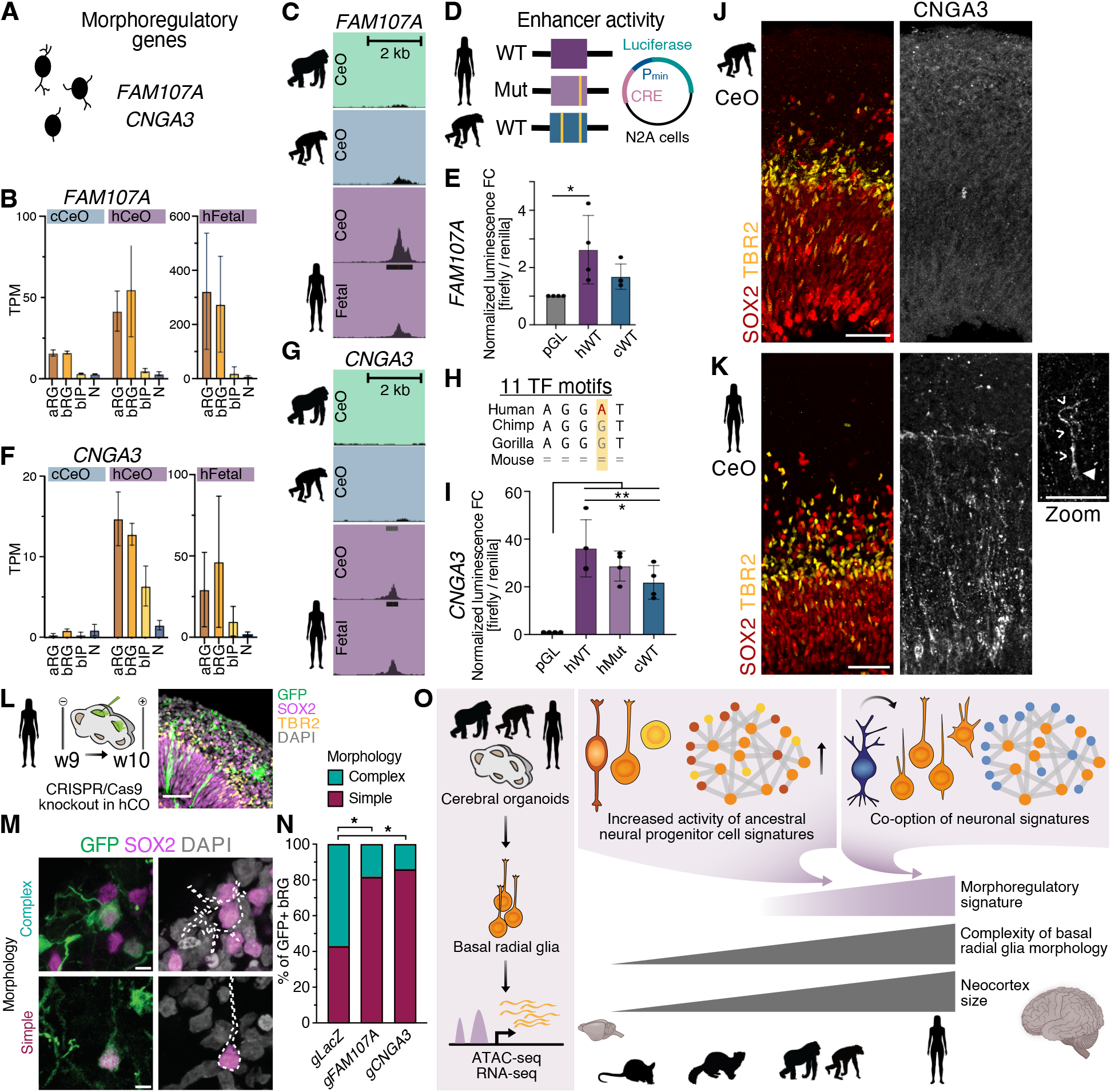
*FAM107A* and *CNGA3* are required for the morphological complexity of human bRG. (**A**) Analysis of morphoregulatory genes. (**B**) Gene expression of *FAM107A* determined by RNA-seq. Mean ± SD displayed in TPM. (**C**) Chromatin accessibility determined by ATAC-seq for the *FAM107A*-linked hbRG cCRE (chr3: 59,505,735–59,506,833). Merged bigWig tracks of at least two replicates, with standardized track height 0–3. Called peaks are represented by black bars in tracks. (**D**) Enhancer activity is determined by luciferase reporter gene assays in mouse N2A cells for the 230 bp-center hbRG cCRE human wildtype sequence, a mutated human sequence with reverted chimpanzee nucleotide in the transcription factor binding site, and the chimpanzee orthologous sequence. (**E**) Normalized luminescence activity for the empty reporter backbone (pGL) and human and chimpanzee wildtype *FAM107A*-linked hbRG cCRE sequences. Mean ± SD from four independent experiments; * p < 0.05, ** p < 0.01; ANOVA with Tukey’s multiple comparison post hoc test. (**F**) Gene expression of *CNGA3*. (**G**) Chromatin accessibility for the *CNGA3*-linked hbRG cCRE (chr2: 98,149,173–98,149,634). (**H**) The sequence of the 230bp *CNGA3*-linked hbRG cCRE peak center contains a human-specific nucleotide change, creating human-specific binding sites for eleven transcription factors. (**I**) Normalized luminescence activity for the *CNGA3*-linked hbRG cCRE sequences. (**J, K**) Immunohistochemistry for SOX2, TBR2, and CNGA3 in chimpanzee (J) and human (K) CeOs shows protein expression in human radial glia, including bRG (zoom), but not chimpanzee. Scale bars, 50 μm. (**L**) Scheme of the CRISPR/Cas9-mediated knockout in hCOs. Electroporated hCO ventricle analyzed by immunofluorescence for GFP, SOX2 and TBR2, and staining for DAPI. Scale bar, 50 μm. (**M**) Examples of electroporated GFP+ SOX2+ TBR2– bRG with simple and complex morphologies in hCOs revealed by immunofluorescence for GFP and SOX2, and staining for DAPI. Scale bars, 5 μm. (**N**) Relative distribution of bRG according to their morphological complexity (*13*). Mean of 3 independent hCO differentiations and KO experiments. * p < 0.05; 2-way ANOVA with Bonferroni post hoc test. (**O**) Summary of findings. Interspecies comparison of bRG isolated from CeOs suggest that human-specific morphoregulatory signatures characterize neocortex evolution. WT, wildtype; Mut, mutated; Pmin, minimal promoter; TPM, transcripts per million; TF, transcription factor.

Another example is CNGA3, a cGMP-activated depolarizing ion channel described to be expressed in cone photoreceptors, whose dysregulation leads to abnormal neurite outgrowth in the retina (*56*). Recent studies describe expression in non-cortical brain regions (*57*). In CeOs, *CNGA3* expression is largely absent in chimpanzees, whereas *CNGA3* is highly expressed in human neural progenitor cells (Fig. 4F). The *CNGA3*-linked hbRG cCRE (Fig. 4G) has no ortholog in mice and contains within its central region a human-specific G-to-A conversion creating putative binding sites for eleven transcription factors (Fig. 4H), including ELF1, previously shown to drive glioma proliferation (*57*) and ELK1, essential for neuronal function and cytoskeletal dynamics (*58*). Luciferase reporter assays show substantial gene regulatory potential for both the human and chimp *CNGA3* cCRE sequences, with the human sequence showing significantly higher activity (Fig. 4I). As with *FAM107A*, the hbRG cCRE targeting *CNGA3* can thus be shown to be a bona fide enhancer with higher activity potential in humans than in chimpanzee. As expression of CNGA3 had so far not been described in the developing neocortex, we investigated protein expression in great ape CeOs. Chimpanzee CeOs did not show protein expression of CNGA3 (Fig. 4J), whereas CNGA3 was clearly detectable in human CeOs, including in cells with typical bRG morphology (Fig. 4K). It thus appears that human radial glia co-opted *CNGA3* expression from ancestral expression in the retina or non-cortical neurons, underlining the concept that altered morphoregulatory dynamics in human bRG are partially based on re-purposing existing CREs and their target genes.

### *FAM107A* and *CNGA3* are required for the morphological complexity of human bRG

To directly assess whether *FAM107A* and *CNGA3* influence human bRG morphology, we performed CRISPR/Cas9-mediated knockout by electroporating human cortical organoids (hCOs) with ribonucleoprotein complexes (Fig. 4L; fig. S5) (*22, 59*). We then analyzed the morphology of SOX2+ TBR2– bRG in the SVZ of hCOs, which contain previously described bRG morphotypes (*13*), including monopolar, bipolar, multipolar, and nonpolar cells, as well as cells with bifurcated processes (Fig. 4M; fig. S6). Morphological analysis reveals that the proportion of bRG with complex morphologies is reduced upon knockout of *FAM107A* and *CNGA3* compared to control (Fig. 4N; fig. S6). Taken together, these results further support the concept that human bRG gene regulatory networks enhance the activity of morphoregulatory pathways, leading to more complex bRG morphology in human cortical development (Fig. 4L).

## Conclusion

In this study, we identified gene-regulatory patterns specific to human bRG that may underly their unique biology, particularly their extensive proliferative potential and neuronal output–both critical for neocortex expansion during evolution (*3, 4, 17*). We discovered a consistent pattern of human bRG gene regulatory regions, their target genes, and human-upregulated genes involved in the regulation of neuronal cell morphology. Neurons are characterized by great morphological complexity that underlies their specific connectivity patterns (*60*). Our data show strongly increased accessibility in ancestral neural progenitor CREs as well as a subset of human bRG CREs exhibiting concomitant and ancestral activity in neurons, reflected in the expression patterns of their target genes. This suggests that human bRG gained activity in NPC-active morphoregulatory CREs and genes, and may have re-purposed neuronal gene regulatory networks to regulate and likely complexify their morphology. Increased morphological complexity and resulting increased proliferative potential of bRG have been shown to correlate with neocortex size (*13*), likely mediated by the increased exposure to proliferative signals from the extra-cellular environment (*61*). Our data suggest that this pattern extends to a human-specific context and that morphoregulatory signatures distinguish human bRG from other great apes. We propose that the co-option of morphoregulatory and neuronal signatures is a key evolutionary mechanism contributing to these changes.

## Acknowledgments

We are grateful to the facilities of the CRTD and Dresden Concept partners for the outstanding support provided, notably K. Neumann and her team at the Stem Cell Engineering Facility, H. Hartmann and her team at the Light Microscopy Facility, A. Gompf and her team at the Flow Cytometry Facility, S. Weiche from histology, and A. Dahl and A. Kränkel at the DRESDEN-concept Genome Center. We acknowledge the Coriell Institute for providing the human WTC11 iPSC line and Svante Pääbo for kindly sharing the SandraA chimpanzee cell line. We thank all members of the Albert laboratory for their support and discussions. Grammarly was used for spelling and grammar checks. ChatGPT was used in the context of data analysis and coding work. The output of these tools was reviewed and approved by the authors.

## Funding

This work was supported by grants from the

German Research Foundation Emmy Noether AL 2231/1-1 (MA, TMS, AK);

Schram Stiftung (MA, ND);

Federal Ministry of Education and Research grant ERA-NET MEPIcephaly 01EW2208 (MA, ND).

## Author contributions

Conceptualization: TMS, MA;

Investigation: TMS, ND, AK, EC, IC;

Resources: IC, JP, MK, CE, RPD, CB, SP, UM, PW, KL, ML;

Formal analysis: TMS, SV, BB;

Visualization: TMS, ND, MA;

Funding acquisition: ML, KL, NK, BB, MA;

Supervision: ML, NK, BB, MA;

Writing – original draft: TMS, MA;

Writing – review & editing: all authors.

## Data and materials availability

De-identified human and great ape ATAC-seq data have been deposited at Zenodo. Processed ATAC-seq and RNA-seq data are available in Supplementary Data Tables S1 to S3. This study did not generate new cell lines or code. Any additional information required to reanalyze the data reported in this paper is available from the corresponding author upon request. Restrictions apply to human sequencing data due to data privacy regulations and ethical permissions.

## Supplementary Materials

Materials and Methods

Figs. S1 to S6

Tables S1 to S2

References Data S1 to S3

