## Supplementary data for "Human-specific morphoregulatory signatures in basal radial glia characterize neocortex evolution"

**The PDF file includes:**

Materials and Methods
Figs. S1 to S6
Tables S1 to S2
References

**Other Supplementary Materials for this manuscript include the following:**

Data S1 to S3

### Materials and Methods

Human fetal brain tissue

Human fetal brain tissue was obtained from two sources. Firstly, the Department of Gynecology and Obstetrics, University Clinic Carl Gustav Carus of the Technische Universität Dresden, following elective pregnancy termination and informed written maternal consent and with approval of the local University Hospital Ethical Review Committee (ethical approval number EK 355092018). Second, human fetal material was provided by the Joint MRC/Wellcome Trust (grant# MR/X008304/1 and 226202/Z/22/Z) Human Developmental Biology Resource (http://hdbr.org). Fetal age was determined as gestation week (GW) 13 or 14 through ultrasound measurements of crown-rump length and other established developmental stage criteria. Due to protection of data privacy, the sex of the human fetuses supplying the tissue cannot be disclosed; however, sex is unlikely to impact this study's outcomes significantly. No instances of health disorders were documented among the fetal tissue samples utilized in this investigation. Tissue dissection was performed in Tyrode's solution (TS), and the tissue was used for experimental purposes within 1–6 hours following the procedure.

Human and primate induced pluripotent stem cell lines

All iPSC lines were grown under standard conditions (5% $CO_2$, 37°C) on Matrigel-coated plates. The human iPSC lines WTC11 (UCSFi001-A; Gladstone Institutes, distributed by Coriell Institutes) (*1*) and CRTDi004-A (*2*), and chimpanzee iPSC line SandraA (*3*) were cultured in mTeSR1, whereas StemFlex medium was used for the gorilla iPSC line GC1 (*4*). The lines WTC11 and GC1 were passaged using ReLeSR. For CRTDi004-A and SandraA, TrypLE Express enzyme and Y-27632 rho kinase inhibitor were used. All experiments involving human iPSCs were performed in accordance with the ethical standards of the institutional and/or national research committee, as well as with the 1964 Helsinki Declaration, and approved by the University Hospital Ethical Review Committee (IRB00001473; IORG0001076; ethical approval number SR-EK-456092021).

Generation of cerebral and cortical organoids

Cerebral organoids were generated with small modifications as described in detail (*5*) from iPSC lines WTC11 and SandraA. Briefly, 9,000 cells were seeded into 96-well Ultra-Low Attachment plates and cultured in mTeSR1 containing 10 µM Y-27632 rho kinase inhibitor. Plates were spun for 3 min at 300x g to settle cells. After 48 hours, medium was changed to mTeSR1 without Y-27632 rho kinase inhibitor. On day 4, the now formed embryoid bodies (EBs) were changed to neural induction medium. EBs were embedded in Matrigel on day 8, transferred to differentiation medium containing B27 supplement without vitamin A and henceforth cultured on an orbital shaker. Starting on day 14, young organoids were changed to, and then maintained in, differentiation medium containing B27 supplement with vitamin A, changed every 2–3 days until collection for experiments. Gorilla cerebral organoids were generated based on previous protocols (*5, 6*), using the STEMdiff Cerebral Organoid Kit. All cerebral organoids were collected for experiments on w8 ± 3 days.

Knockout experiments were carried out in sliced cortical organoids (*7*) generated using the CRTDi004-A iPSC line, as described before (*8, 9*). In short, iPSC colonies were grown to a diameter of 1.5 mm and detached using collagenase for 1 hour at 37°C to initiate EB formation in forebrain medium 1 in 6-well ultra-low attachment plates. Medium was changed on day 3 and 4 of organoid culture. Subsequently, half of the medium was replaced by forebrain medium 2 at day

5 and 6, before EBs were embedded in Matrigel cookies on day 7 in forebrain medium 2. After one week, cortical organoids were mechanically released from Matrigel and cultured in forebrain medium 3 on an orbital shaker at 120 rpm from then on. On day 35, forebrain medium 3 was supplemented with Matrigel for the remaining culture. Cortical organoids were sliced into 500 µm-thick slices using a vibratome at 45 days of culture. Medium was changed to forebrain medium 4 on day 63.

Mouse Neuro-2A (N2A) cell line
The mouse neuroblastoma cell line was grown in standard conditions (5% $CO_2$, 37°C) on 6-well Nunclon Delta Surface plates. Cells were cultured in DMEM/F12 supplemented with 10% Fetal Bovine Serum and 1% Penicillin/Streptomycin. Cells were passaged using Trypsin-EDTA.

Tissue labeling with the lipophilic dye DiI
The protocol for DiI-labeling was adapted from a previously published study (*10*). For each experiment, about two-thirds of the experimental tissue was labeled in pre-warmed (37°C) TS with CellTracker CM-DiI Dye (1 mM in DMSO), the fixable variant of the previously published DiI for cell sorting (*10*). More specifically, human fetal cortical tissue, roughly amounting to 30–50 million live cells, was labeled by lightly brushing a shortened microloader pipette tip over the entire basal surface of the cerebral hemisphere. After one passage, the hemisphere was transferred to a fresh TS-containing petri dish to wash away excess dye. The labeling and washing process was repeated up to 5 times until the entire basal surface was uniformly coated in CM-DiI, established by inspection under a fluorescent microscope. After labeling, primary tissue was transferred to 20 mL glass flasks containing DMEM/F12 supplemented with 1x N2 and 1x B27 with 1–3 cortical fragments per flask. Tissue was incubated under rotation (FFTC) (*11*) with 40% $O_2$, 5% $CO_2$ and 55% $N_2$ at 37°C in a whole-embryo culture incubator for 4 hours up to overnight. For cerebral organoids, the entire surface of 5–10 organoids was labeled, and organoids subsequently returned to culture in their respective medium for 3 hours. For all experiments, unlabeled tissue was cultured separately but under the same conditions as labeled tissue. Post-incubation, organoids were further dissected in pre-warmed (37°C) TS to enrich for visible ventricle-like regions.

Immunocytochemistry
Experimental tissue was dissociated using the MACS Neural Tissue Dissociation Kit P with minor modifications for each tissue. For organoids, 3–6 organoids, depending on size, were combined in 1 mL enzyme mix 1 (50 µL enzyme P + 950 µL buffer X), incubated for 6 min at 300 rpm and at 37°C in a thermoshaker, quenched with 30 µL enzyme mix 2 (10 µL enzyme A + 20 µL buffer Y) and conditions recombined after trituration to single cells. For fetal tissue, up to 400 mg of tissue was combined in 1950 µL of enzyme mix 1 (50 µL enzyme P + 1900 µL buffer X), incubated for 15 min, quenched with 30 µL enzyme mix 2 (10 µL enzyme A + 20 µL buffer Y) and, if applicable, conditions recombined after trituration to single cells. After trituration to single cells, all tissues were spun for 5 min at 300x g and 4°C to pellet cells and re-suspended in 1 mL cold TS. Cells were counted, and viability was assessed using 10 µL of the cell suspension by Trypan blue staining.
The protocol for immunocytochemistry, subsequent cell sorting and processing was adapted from previously established approaches (*12, 13*). All centrifugation steps were carried out at 4°C unless otherwise noted. For fixation, cells were re-suspended to no more than 20 million cells/mL and

incubated at 1% v/v formaldehyde in TS for exactly 10 min at 10 rpm rotation at room temperature (RT). Fixation was quenched with 0.2 M glycine for 5 min at 10 rpm rotation. Cells were pelleted for 5 min at 500x g, washed once in 1% BSA in TS and re-pelleted the same way. Cells were permeabilized on a rocking platform for 10 min at 4°C in buffer containing 0.1% saponin and 0.2% BSA in TS and subsequently pelleted for 3 min at 2,500x g. For antibody staining, 4 million cells each were aliquoted from the unlabeled tissue condition for single stain controls as well as an unstained control. From the DiI-labeled tissue condition, 4 million cells were aliquoted for a DiI-only control. Non-control conditions were continued in the presence of 1:1,000 RNAsin Plus. Antibody staining was performed in the dark for 1 hour on a rocking platform at 4°C in buffer containing 0.1% saponin and 1% BSA in TS as follows: Human and primate DiI-labeled condition: V450 Mouse anti-Sox2 (1:40) or Rabbit anti-Pax6 (1:350), FITC Rat anti-Ctip2 (1:500) or Rat anti-Ctip2 (1:500) and Alexa Fluor 647 Mouse anti-ZO-1 (1:750). Human and primate unlabeled condition: V450 Mouse anti-Sox2 (1:40) or Rabbit anti-Pax6 (1:350), FITC Rat anti-Ctip2 (1:500) or Rat anti-Ctip2 (1:500), Alexa Fluor 647 Mouse anti-ZO-1 (1:750) and PE Mouse anti-EOMES (1:33). Subsequently, cells were pelleted for 3 min at 2,500x g and washed twice in buffer containing 0.1% saponin and 0.2% BSA in TS. Conditions stained with Rabbit anti-Pax6 or Rat anti-Ctip2 were subsequently stained with Goat anti-Rabbit Alexa Fluor 405 or Donkey anti-Rat Alexa Fluor 488 in the dark for 1 hour on a rocking platform at 4°C in buffer containing 0.1% saponin and 1% BSA in TS and then washed twice in buffer containing 0.1% saponin and 0.2% BSA in TS. All cells were re-suspended to 1–3 million cells/100 µL in 0.5–1 mL 0.5% BSA and 1:1,000 RNAsin Plus in TS, passed through the 35 µm filter of their respective test tube and kept on ice for flow cytometry analysis.

Fluorescence activated cell sorting

Cells were sorted with a 100 µm nozzle with a sample chamber and tube holder at 4°C. For each experiment, drop delay was determined, and fluorophore compensation was calculated based on single-stain and unstained conditions. The main cell populations were defined using side scatter (SSC) area and forward scatter (FSC) area. Cell debris and aggregates are excluded using FSC height against FSC-A and SSC width against SSC-H. Cells were sorted using a BD FACSAria II with BD FACSDiva Software (v8.0.2 or v9.0.1). Cell populations were collected in DNA LoBind tubes containing either ATAC-RSB (*14*) for subsequent ATAC-seq, or a mix of 95 µL RNAse-free $H_2O$, 95 µL 2x Digestion Buffer and 10 µL Proteinase K (Zymo QuickRNA FFPE MiniPrep) for subsequent RNA isolation, and kept on ice until further processing.

For each experiment, 50,000 cells of each cell type were sorted for ATAC-seq, 10,000 or 5,000 cells were sorted for RNA-seq and 5,000 cells were sorted for analysis by RT-qPCR. Deep-layer neurons (DL-N) and basal intermediate progenitor cells (bIPs) were isolated from cell suspensions without DiI label. DL-N were defined as SOX2–/CTIP2+/ZO1–/TBR2– and gated in this hierarchy. bIPs were defined as not restricted on CTIP2 expression (CTIP2±) and gated for TBR2+/SOX2–/ZO1– in this hierarchy. Apical and basal radial glia were isolated from DiI-labeled cell suspensions. aRG were defined as SOX2+/ZO-1+/DiI+, while bRG were defined as SOX2+/ZO-1–/DiI+ and gated in this hierarchy. CTIP2 staining was used as an internal biological control but not gated, as radial glia target cells were always CTIP2–. SOX2 and PAX6 antibodies were used interchangeably.

ATAC-seq library preparation

ATAC-seq was performed as previously described with minor alterations to the protocol (*14*). Briefly, sorted cells are pelleted at 500x g and 4°C for 5 min. All supernatant was aspirated, cells were re-suspended in 50 µL ice-cold lysis buffer by pipetting up-and-down three times and incubated on ice for 3 min. Lysis was washed out with 1 mL ice-cold wash buffer and tubes inverted to mix. Cells were again pelleted at 500x g and 4°C for 10 min. Supernatant was fully aspirated and cells re-suspended in 50 µL transposition mix containing 100 mM transposase. Reactions were incubated at 37°C and 1,000 rpm in a thermoshaker for 2 hours and then quenched by addition of 40 mM EDTA. Reactions were pelleted at 500x g for 5 min, re-suspended in 200 µL reverse-crosslink solution and incubated O/N at 65°C and 1,000 rpm in a thermoshaker as previously reported for ATAC-seq on fixed cells (*15*). The next day, the original protocol (*14*) was continued, and reactions were cleaned up using a Zymo DNA Clean and Concentrator-5 Kit, libraries were amplified with determination of amplification cycles by qPCR as described (*14*) and unique adapter combinations (i5 and i7), and final PCR reactions were cleaned up using a Zymo DNA Clean and Concentrator-5 Kit. Final libraries were size selected using SPRIselect Beads at 0.5x and 1.3x ratios, to retain fragments of 150–1,000 bp. Successful transposition was established by presence of characteristic periodic nucleosomal pattern on a FragmentAnalyzer NGS-Kit 1-6000bp (Agilent), libraries subsequently quantified using a Qubit fluorometer (Invitrogen) and at least 50 million fragments were sequenced with 100 bp paired-end reads on an S4 v.15 200 cycles flow cell on the NovaSeq 6000 machine (Illumina).

RNA-seq library preparation

RNA extraction was performed as per manufacturer's instructions using the Zymo QuickRNA FFPE MiniPrep Kit and columns were eluded with 10 µL DNase/RNase-free $H_2O$ and stored at -80°C until library preparation. Transcriptome libraries were prepared using the SmartSeq 2 protocol, modified from (*16*). Isolated total RNA from 10,000 cells was denatured for 3 min at 72°C in 4 µL hypotonic buffer (0.2 % Triton X-100) in the presence of 2.4 mM dNTPs, 240 nM dT-primer and 4 U RNase Inhibitor (NEB). Reverse transcription was performed at 42°C for 90 min after filling up to 10 µL with RT buffer mix for a final concentration of 1x Superscript II buffer, 1 M betaine, 5 mM DTT, 6 mM $MgCl_2$, 1 µM TSO-primer, 9 U RNase inhibitor and 90 U Superscript II Reverse Transcriptase. The reverse transcriptase was inactivated at 70°C for 15 min. For subsequent PCR amplification of the cDNA, the optimal PCR cycle number was determined with an aliquot of 1 µL unpurified cDNA in a 10 µL qPCR containing 1x KAPA HiFi Hotstart Readymix, 1x SYBR Green I and 0.2 µM UP primer. The determined cycle number was then used to amplify the residual 9 µL cDNA using 1x KAPA HiFi HotStart Readymix and 250 nM UP-primer under the following cycling conditions: initial denaturation at 98°C for 3 min, 12–17 cycles [98°C 20 sec, 67°C 15 sec, 72°C 6 min] and final elongation at 72°C for 5 min. The amplified cDNA was purified using 0.6x Sera-Mag SpeedBeads and resuspended in a buffer consisting of 10 mM Tris, 20 mM EDTA, 18.5 % (w/v) PEG 8000 and 2 M sodium chloride solution. cDNA quality and concentration was determined using the FragmentAnalyzer NGS 1-6000bp Kit.

For library preparation, 2 µL amplified cDNA was tagmented in 1x Tagmentation Buffer using 0.8 µL bead-linked transposome (Illumina) at 55°C for 15 min in a total volume of 4 µL. The reaction was stopped by addition of 1 µL 0.1 % SDS (37°C, 15 min), beads were bound to a magnet and the supernatant was removed. Beads were then resuspended in 14 µL indexing PCR Mix containing 1x KAPA Hifi HotStart Ready Mix and 700 nM unique dual indexing primers (i5 and i7), and subjected to a PCR (72°C 3 min, 98°C 30 sec, 12 cycles [98°C 10 sec, 63°C 20 sec, 72°C 1 min], 72°C 5 min). Libraries were purified with 0.9x volume Sera-Mag SpeedBeads, followed by a double size selection with 0.6x and 0.9x. Repurification with a final 0.9x bead purification was performed if needed. Sequencing was performed after quantification using the FragmentAnalyzer NGS 1-6000bp Kit (Agilent) on a NovaSeq S4 v1.5 200 cycles flow cell (Illumina) to a sequencing depth of at least 50 million 100 bp paired-end reads per library.

Gene expression analysis by RT-qPCR

RNA extraction was performed as per manufacturer's instructions using the Zymo QuickRNA FFPE MiniPrep Kit. cDNA was synthesized using random hexamers and Superscript III Reverse Transcriptase. Final products were diluted by addition of 180 µL $H_2O$ and stored at -20°C until further analysis by RT-qPCR. qPCR was performed in technical triplicate using LightCycler 480 SYBR Green I Master on a Light Cycler 480 (Roche). Data was normalized by *GAPDH* expression. Primers are listed in Table S1.

Cloning

The center 230 bp of candidate regions were synthesized for the human, human-mutated and chimpanzee sequences with restriction sites (IDT gBlocks Gene Fragments) for KpnI and BmtI. Fragments were inserted into the pGL4.23[luc2/minP] backbone (Promega) using NEB KpnI-HF and NEB BamHI-HF with CutSmart buffer according to manufacturer's instructions and confirmed by Sanger sequencing. Sequences are listed in Table S1.

Luciferase assay

N2A cells were seeded at a density of 10,000 cells/well in a white-walled, clear bottom 96-well plate one day before lipofection. Lipofection was performed according to manufacturer's instructions using Lipofectamine 2000 DNA Transfection Reagent (Invitrogen) with 2.5 µL of lipofectamine, 50 ng pRL-CMV (Promega) vector for normalization and 450 ng of pGL4.23[luc2/minP]-construct vector.
Luciferase assays were performed as previously described (*17*). 24 hours after lipofection, Dual-Luciferase Reporter Assay System from Promega was used according to manufacturer's instructions in concert with EnVision Multimode plate reader by PerkinElmer to quantify CRE activity of candidate regions. All experiments were performed in triplicate wells/construct and 3 technical measurements/well were acquired for both Firefly (pGL4.23[luc2/minP]-construct vector) and Renilla (pRL-CMV). Raw Firefly measures were normalized to Renilla activity for each well. The experiment was independently repeated four times.
The arithmetic means of Firefly/Renilla triplicate technical measurements in each well were calculated and subsequently, the values from all three wells/construct averaged for each experiment. Shapiro-Wilk and Levene's test were used to assess normality and homoscedasticity, respectively, followed by one-way ANOVA and Tukey's post-hoc. The cut-off for significance was set to $p < 0.05$.

ATAC-seq data analysis

ATAC-seq libraries were mapped and processed using the ENCODE pipeline (*18*) with default settings. GorGor6, panTro6 and hg38 were used as the reference genomes for gorilla, chimpanzee and human, respectively. Putative orthologous regulatory regions were identified using the HALPER tool (*19, 20*) after generating reference-free Cactus multiple sequence alignments of the gorGor6, panTro6 and hg38 genomes (*21*). The orthologous regions were merged to generate a common peak set (225,395 peaks) with the genomic coordinates of each corresponding species.

Distal chromatin peaks refer to all peaks that do not overlap a transcription start site. DESeq2 (*22*) was used to calculate differentially accessible regions based on species (gorilla, chimpanzee and human) and cell type (aRG, bRG, bIP and N). Differentially accessible regions were intersected with genome coordinates of previously assembled candidate cis-regulatory elements (*23*) and human-accelerated regions (*24*). cCRE were linked to their target genes by generating all possible pairs with transcription start sites (TSS). Distal peaks were associated with genes located within the same topologically associating domain (TAD) that had the highest Hi-C score, provided the distance ranged between 5 kb and 2 Mb. These analyses were performed using previously generated published Hi-C data from organoid RGs and IPs (*25*). GO term analysis (*26, 27*) on target genes was performed using the packages 'enrichr' (v3.2), 'org.Hs.eg.db' (v3.19.1) and 'clusterProfiler' (v4.12.6).

Transcription factor motif analysis

The JASPAR2020 core vertebrate database was used for motif-based analysis. Transcription factor motif enrichment was calculated using the monaLisa package (v1.10.0) (*28*). 'Motifmatchr' (v1.26.0) was then used to identify transcription factor motifs within genomic regions (p.cutoff = 0.0005) and center the region around them.

RNA-seq data analysis

RNA-seq libraries were mapped and de-duplicated using STAR (*29*) with default settings. Hg38 was used as the reference genome for both human and chimpanzee (*30*). All downstream analysis was conducted in R v4.4.1. DESeq2 (*22*) was used to calculate FPKM and differentially expressed genes (FDR < 0.05, log2FC = 1) based on species (chimpanzee and human) and cell type (aRG, bRG, bIP, and N). GO term analysis (*26, 27*) was performed using the packages 'enrichr' (v3.2), 'org.Hs.eg.db' (v3.19.1) and 'clusterProfiler' (v4.12.6). The morphoregulatory pathway activity score was adapted from a previous publication (*31*) using the GO terms 'GOBP Cell morphogenesis.v2024.1', 'GOBP Cell projection morphogenesis.v2024.1' and 'GOBP Cell projection organization.v2024.1. For TPM values, RNA-seq libraries were aligned to GRCh38 using kallisto (v0.64.1) (*32*) after quality control using FastQC (v0.11.6). TPM values were obtained in R using 'Tximport' (*33*) using the input 'lengthScaledTPM' together with the 'EnsDb.Hsapiens.v86' package.

Electroporation of cortical organoids

CRISPR/Cas9 gene ablation in human cortical organoids was carried out as described before (*8, 9*). Briefly, cortical organoids were sliced to 500 µm thickness two days before electroporation using a vibratome. On the day of electroporation (day 63 of hCO culture), ribonucleoproteins (RNP) were assembled freshly and separately per guide RNA (gRNA). Two gRNA each targeting either *FAM107A* or *CNGA3* were used, as well as one gRNA targeting *LacZ* as control (sequences are listed in Table S1). Both RNP complexes targeting the same gene were mixed at equimolar ratio and 225 ng/µL pCAG-GFP was added, as well as 0.1 % FastGreen in $H_2O$. Organoids were collected in pre-warmed TS and outwards facing ventricle-like structures were injected with the prepared mixture. Injected organoids were transferred into an electroporation chamber in TS and electroporation was carried out with five pulses applied at 38 V for 50 ms each at intervals of 1 s. Afterwards, electroporated hCOs were transferred into forebrain medium 4 (*7*) and cultured on an orbital shaker for 7 days before immunofluorescent analysis.

CRISPR/Cas9-mediated gene knockout

Guide RNA design and validation for CRISPR/Cas9 gene knockout was performed as previously described (*8, 9*). First, each gRNA was tested separately *in vitro* using PCR products spanning the target region in the genes of interest. After digesting the PCR products with the RNP, fragments were analyzed by gel electrophoresis. Second, the two guide RNAs targeting the same gene were nucleofected together with a pCAG-GFP plasmid into the same iPSC line that was also used for the generation of human cortical organoids, namely CRTDi004-A, with the P3 Primary Cell 4D Nucleofector X Kit S. After 3 days of culture, cells were sorted for GFP by FACS and genomic DNA was extracted. Targeted regions were amplified by PCR, purified after gel electrophoresis and analyzed by Sanger sequencing.

Immunohistochemistry and imaging

All tissue was fixed in 4% paraformaldehyde in 120 mM phosphate buffer pH 7.4 at 4°C for 30 min (organoids) or O/N (human primary tissue). Tissue was transferred to 15% sucrose for 24 hours (human primary tissue only), transferred to 30% sucrose for at least 24 hours, embedded in 50:50 O.C.T. compound : 30 % sucrose in PBS and cut into 20 µm (organoid) or 12 µm (human primary) sections using a cryostat. For immunofluorescence, antigen retrieval was performed in 10 mM citrate buffer pH 6.0 for 1 hour at 70°C and washed out three times with PBS. Sections were quenched in 0.1 M glycine in PBS at RT for 30 min and again washed in PBS. Sections were permeabilized for 30 min at RT in 0.1% Triton X-100 and 10% horse serum and subsequently incubated with primary antibodies in the same buffer O/N (Table S2). After 3 washes with PBS, sections were incubated with secondary antibodies and DAPI (all 1:1,000) for 1 hour at RT in 0.1% Triton X-100 and 10% horse serum before washing out with PBS and mounting in Mowiol. Images were acquired with a Zeiss ApoTome2 fluorescence microscope using a 10x or 20x objective and 1.5 µm thick optical sections, a Zeiss LSM 780/FCS microscope using a 20x, 40x (water immersion) or 63 x (oil immersion) objective, or a Zeiss LSM 980/MP microscope using a 63 x (oil immersion) objective and 0.3 µm thick optical sections. When images were taken as tile scans, the stitching of tiles was performed using the ZEN software.

Analysis of bRG morphology

Morphological analysis of bRG was performed as established earlier (*34*). Individual GFP+ SOX2+ TBR2– bRG in the electroporated organoid SVZ were isolated by cropping an area of 53.3 µm x 53.3 µm surrounding the cell. Cell morphology was analyzed through a Z-stack using GFP signal as a reference to cell shape and assigned to a particular morphotype according to the number and the orientation of processes with respect to the cell body and the VZ, defined as monopolar, bipolar, bifurcated, multipolar or nonpolar. Bonferroni post-hoc test values for Fig. 4N comparing simple and complex morphotypes in g*LacZ* vs. g*CNGA3* *p* value = 0.0037, and g*LacZ* vs. g*FAM107A* *p* value = 0.0085. For Fig. S6B, monopolar g*LacZ* vs. g*CNGA3* *p* value = 0.0159; multipolar *p* value = 0.0126; monopolar g*LacZ* vs. g*FAM107A* p value = 0.0395; multipolar *p* value = 0.0071.

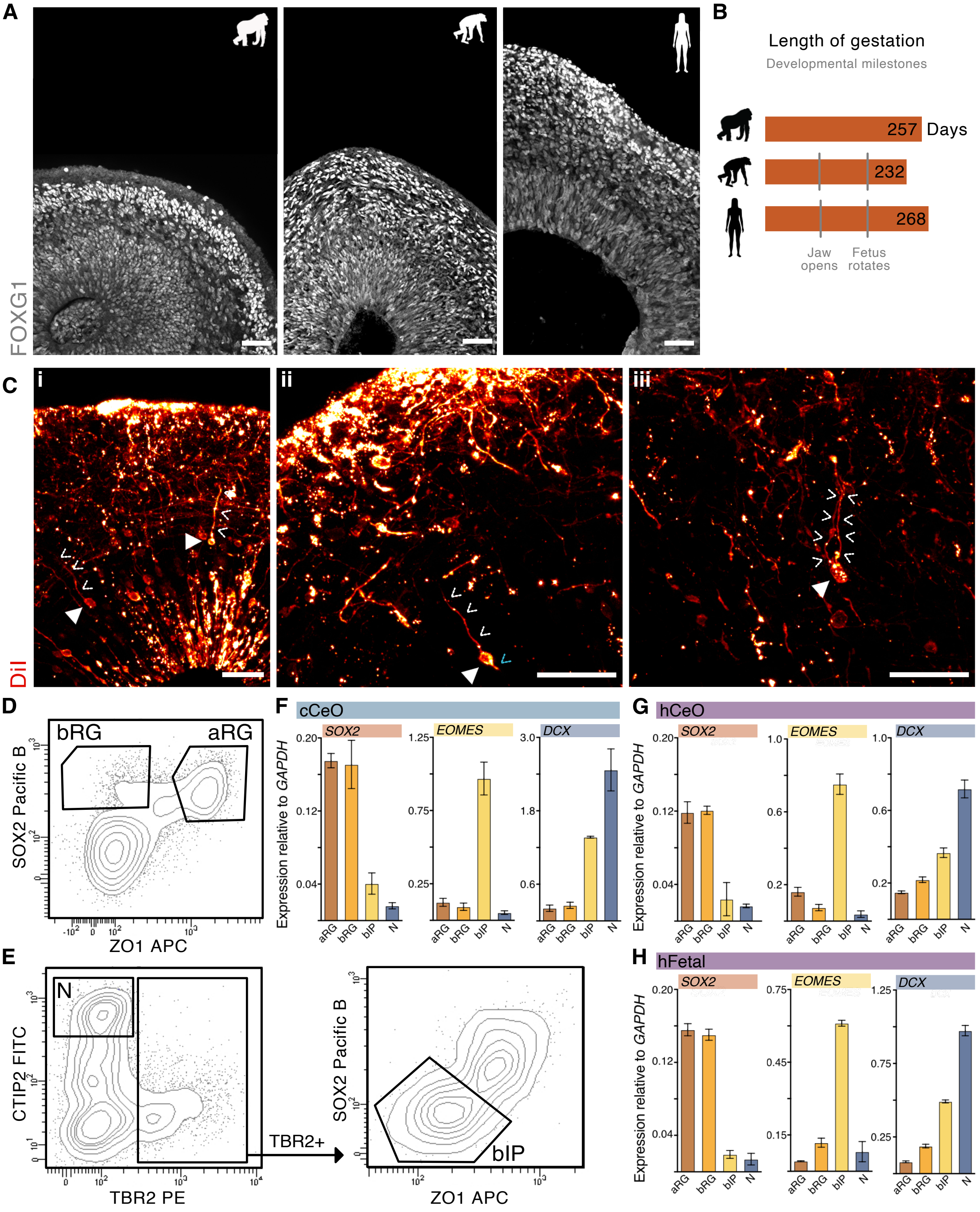


**Fig. S1.**

**Characterization of great ape cerebral organoids and isolated cell types.**

(**A**) CeOs of gorilla, chimpanzee and human at w8 express the forebrain marker FOXG1 denoting cortical identity. Scale bars, 50 µm. (**B**) Comparison of gestation length and timing of *in utero* developmental milestones between gorillas, chimpanzees, and humans indicates that early developmental events are highly correlated in timing (*35, 36*). (**C**) Human CeOs at w8 labeled with the lipophilic dye DiI reveal cells with typical bRG morphology, including previously described morphotypes (*34*) with (**i**) a single basal process, (**ii**) a basal and an apical process, and (**iii**) a bifurcated basal process. Filled arrowheads denote bRG cell bodies, white open arrowheads bRG basal processes and blue open arrowheads bRG apical processes. Scale bars, 50 µm. (**D**) FACS plots of GW13 human fetal cortex showing separation of aRG and bRG based on expression of ZO1, whereas SOX2 is expressed in both RG populations. Both cell populations are further sub-gated for DiI to select cells with basal processes. (**E**) FACS plot of the same GW13/14 human fetal cortex sample, showing gating for neurons as CTIP2+/TBR2– and gating for bIPs as TBR2+/SOX2–/ZO1–. (**F–H**) RT-qPCR of the canonical RG marker gene *SOX2*, the bIP marker gene *EOMES*, and the neuronal gene *DCX*, normalized to the housekeeping gene *GAPDH*, in sorted cell populations of cCeO (F), hCeO (G) and hFetal (H) confirms cell identity on RNA level. Error bars represent standard deviation.

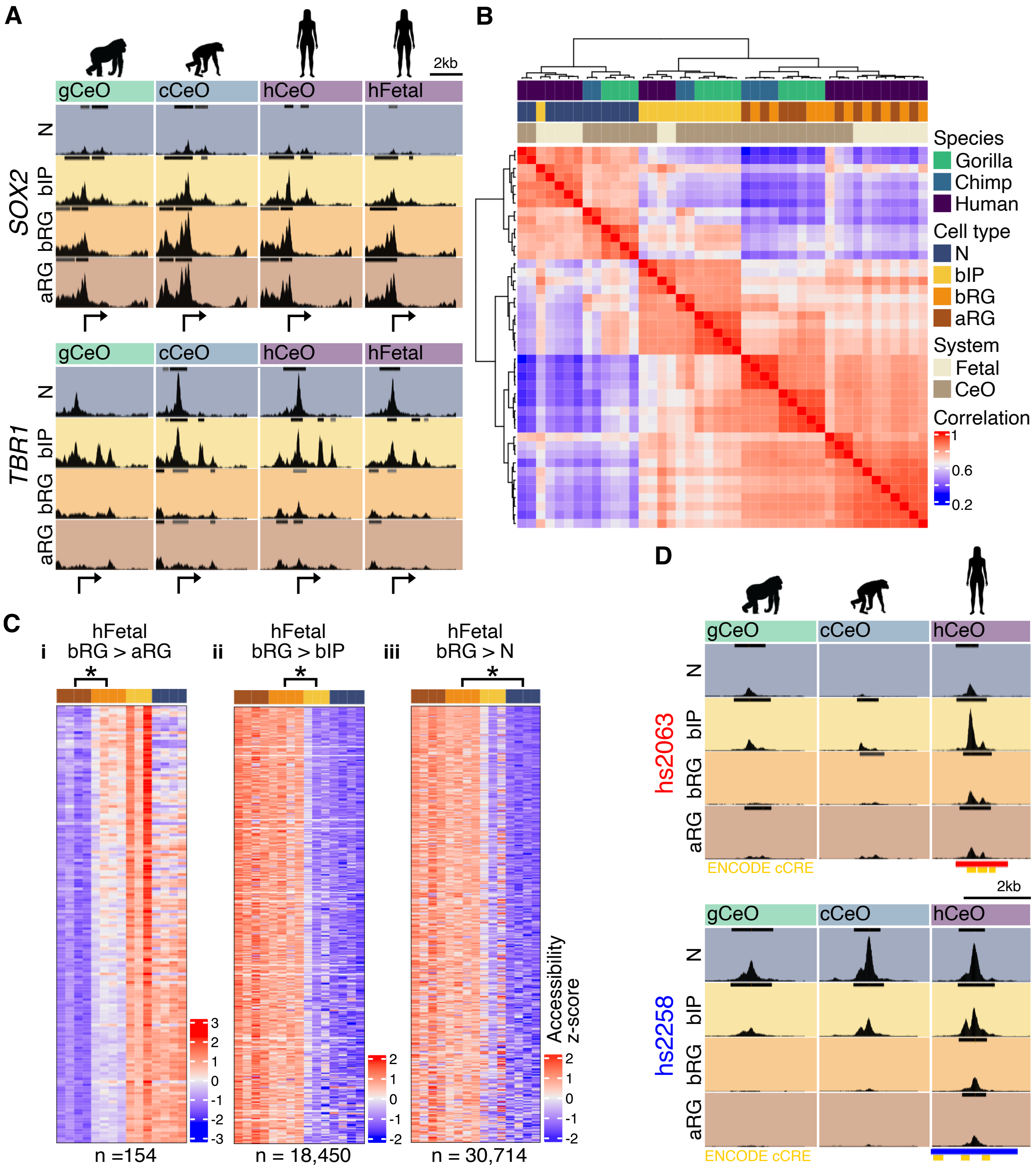


**Fig. S2.**

**Characterization of chromatin accessibility patterns across cell types and species.**
(**A**) Cell type-specific chromatin accessibility exemplified at the transcription start sites (arrows) of the radial glia gene *SOX2* (top) and early neuronal gene *TBR1* (bottom) for each species. Shown are merged bigWig tracks, with standardized track height 0–4.5. Called peaks are represented by black bars in tracks. (**B**) Correlation heatmap of chromatin accessibility for all called peaks across all tissues and cell types indicating patterns of cell type- and species-specific accessibility. (**C**) Chromatin accessibility patterns across all cell types of human fetal cortex tissue of differentially

accessible regions (log2FC ≥ 1; p ≤ 0.05), comparing (**i**) aRG vs. bRG, (**ii**) bIP vs. bRG, and (**iii**) N vs. bRG, underlining that bRG possess a unique molecular profile of sharing variable accessibility features with other cell types rather than being defined by 'bRG-specific' chromatin accessibility. Accessibility z-scores shown for all samples. (**D**) Cell type-specific chromatin accessibility at two characterized VISTA enhancers (*37*), hs2063 (red; top) and hs2258 (blue; bottom), also marked by ENCODE cCRE signatures (yellow) (*23*). Increased activity in human NPCs compared to gorilla and chimpanzee NPCs can be observed in both regions. hs2063 was previously shown to drive reporter gene expression in the central nervous system of the mouse at embryonic day 11.5, while hs2258 did not show activity at this timepoint. Shown are merged bigWig tracks, with standardized track height 0–4.5. Called peaks are represented by black bars in tracks.

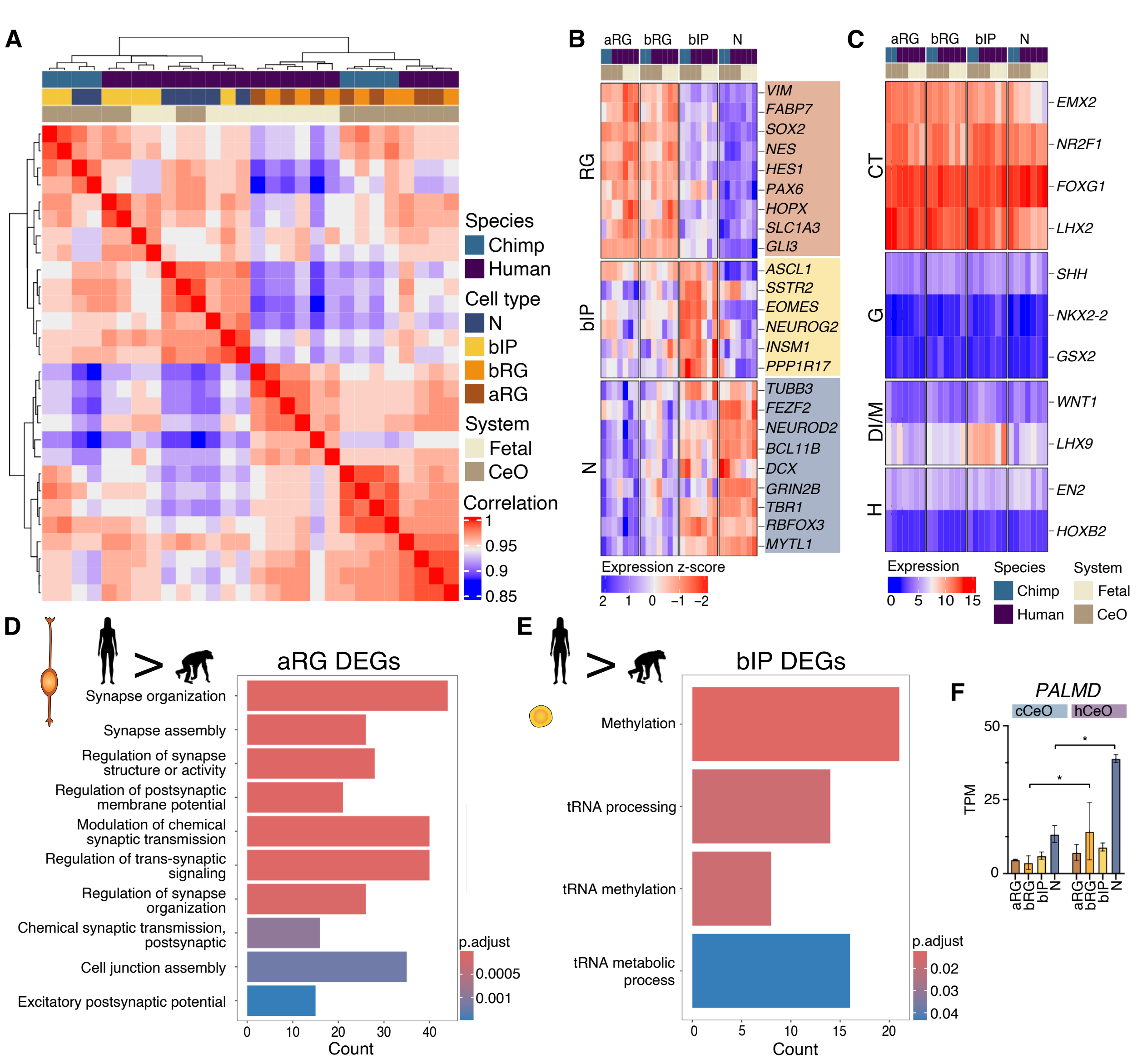


**Fig. S3.**

**RNA-seq reveals cell type- and species-specific expression patterns and confirms tissue identity.**

(**A**) Correlation heatmap of all gene expression patterns reveals cell type and species-specific expression patterns. (**B**) Expression z-score of canonical RG, bIP and neuronal marker genes confirms cell type identity. (**C**) Absolute expression of brain region-specific genes across chimpanzee and human CeOs and human fetal tissue confirm tissue identity at the transcriptional level. (**D**) GO term analysis of genes expressed at higher levels in human vs. chimpanzee aRG indicates enrichment in terms related to regulating synapse structure, but not those regulating other cell processes/protrusions seen in human bRG upregulated genes. Top 10 terms displayed with $p \leq 0.05$. (**E**) GO terms of genes expressed at higher levels in human vs. chimpanzee bIP categorize as regulating several processes related to transfer RNAs. All significant terms displayed with $p \leq 0.05$. (**F**) Gene expression of *PALMD* determined by RNA-seq. Mean ± SD displayed in TPM. *, DESeq2, logFC ≥ 1.5; $p \leq 0.05$. RG, radial glia; bIP, basal intermediate progenitor; N, neuron;

CTX, cortex; GE, ganglionic eminence; DI/MB, diencephalon/midbrain; HB, hindbrain; DEG, differentially expressed gene.

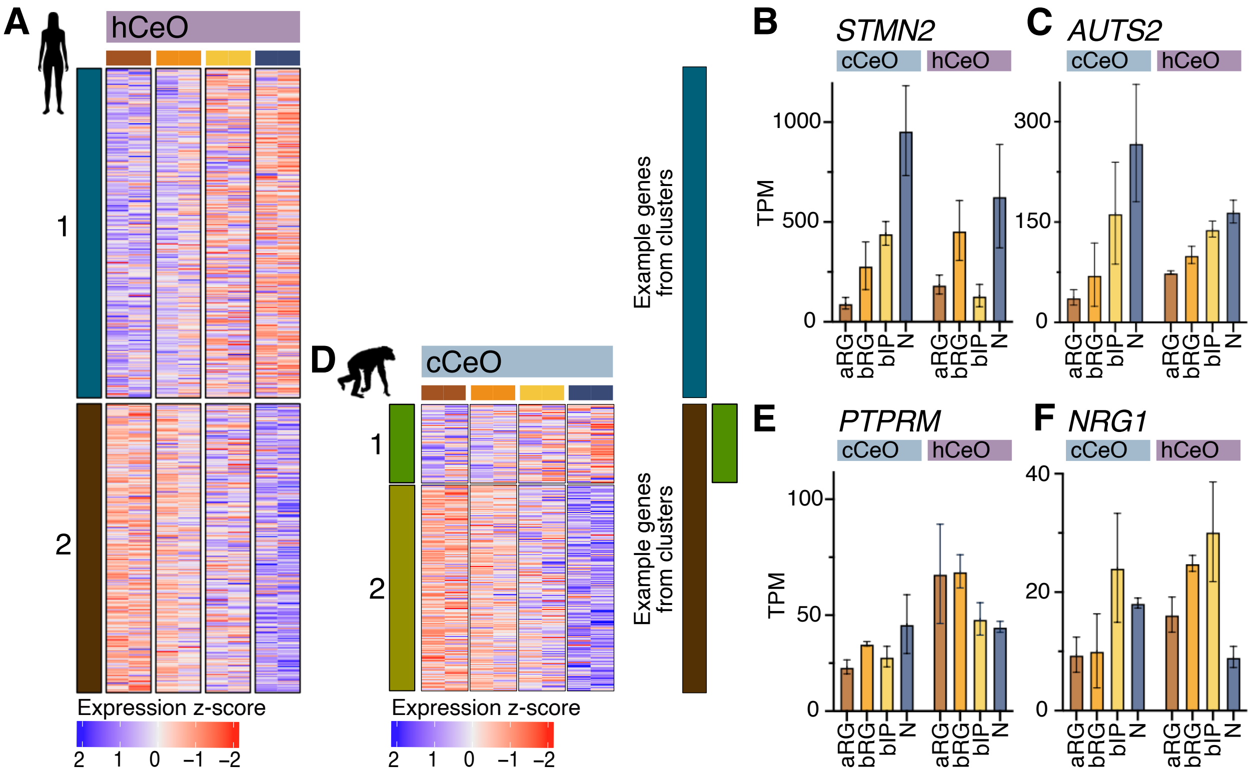


**Fig. S4.**

**Clustering of hbRG cCRE target genes by gene expression.**

(**A**) RNA expression of hbRG cCRE target genes in hCeO reveals two clusters of expression (k-means) across all cell types. Z-score is shown. (**B**) *STMN2* and (**C**) *AUTS2* are two examples of genes in cluster 1, showing higher expression in neurons than neural progenitor cells. Mean ± SD displayed in TPM. (**D**) RNA expression of cluster 2 hbRG cCRE target genes in cCEO shows broadly conserved expression patterns but reveals a subset of genes more highly expressed in bIP and neurons (cluster 1). Z-score is shown. (**E**) *PTPRM* and (**F**) *NRG1* are two examples of genes in human cluster 2 and chimpanzee cluster 1 showing ancestral neuronal expression and significantly increased expression in human neural progenitor cell types. TPM, transcripts per million.

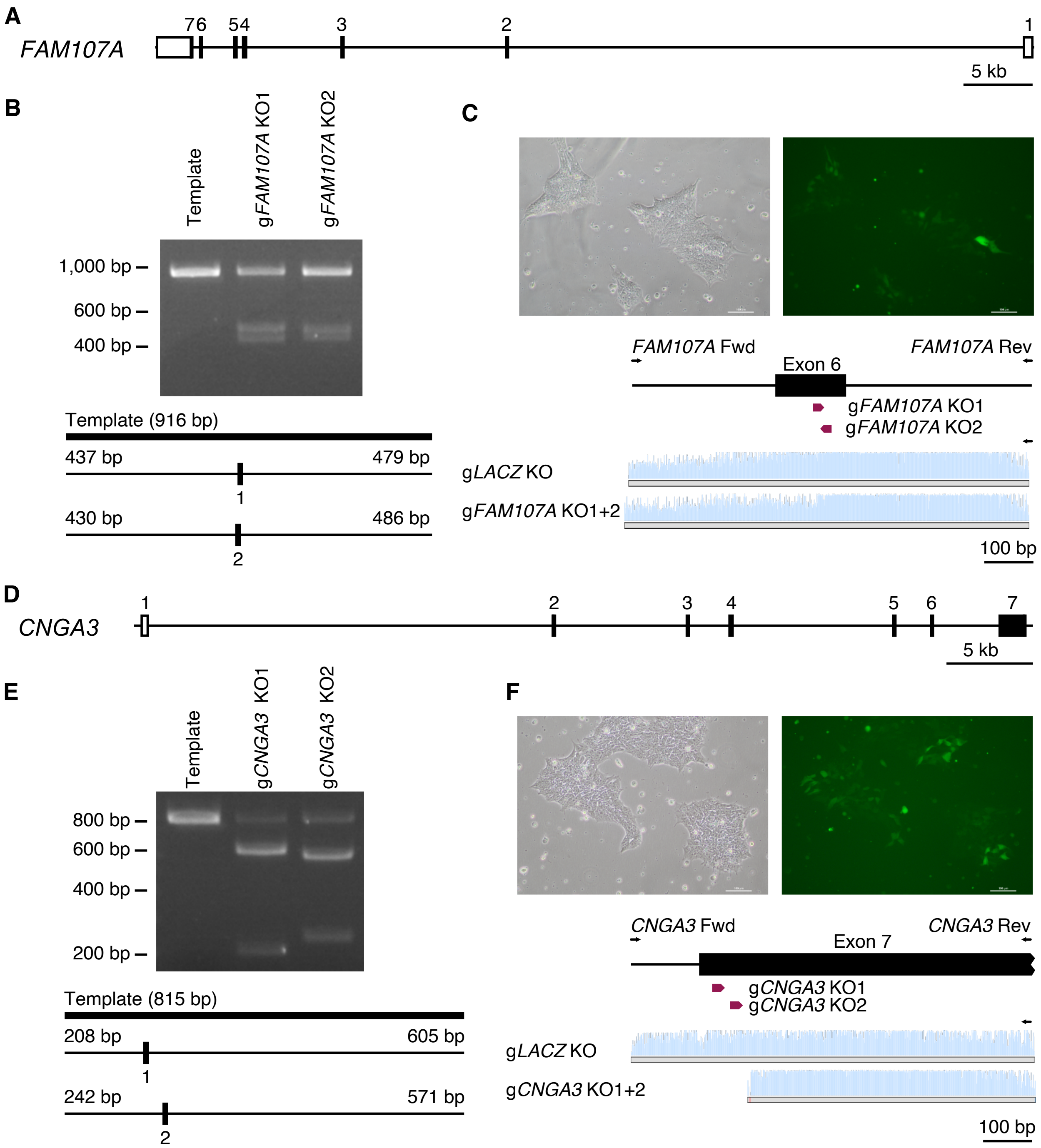


**Fig. S5.**

**CRISPR/Cas9-mediated knockout of *FAM107A* and *CNGA3*.**

(**A**) Schematic illustration of the human *FAM107A* gene locus. (**B**) Guide RNA efficiencies were tested *in vitro*. The effects of the g*FAM107A* KO1+2 RNAs to direct Cas9-mediated cutting of PCR templates was analyzed by agarose gel electrophoresis. Schemes of the sizes of PCR templates, guide RNA binding sites and expected sizes of cut fragments are indicated below. (**C**) CRISPR/Cas9-mediated targeting of *FAM107A* was confirmed in the CRTDi004-A iPSC line by electroporation of Cas9/gRNA ribonucleoprotein complexes together with a GFP plasmid, followed by FACS of GFP-positive cells, PCR amplification of the target region and Sanger sequencing. The sequencing results are shown for g*FAM107A* KO1+2. (**D**) Schematic illustration of the human *CNGA3* gene locus. (**E**) The effects of the g*CNGA3* KO1+2 RNAs to direct Cas9-mediated cutting of PCR templates was analyzed by agarose gel electrophoresis. (**F**) CRISPR/Cas9-mediated targeting of *CNGA3* was confirmed in iPSC. The sequencing results are shown for g*CNGA3* KO1+2.

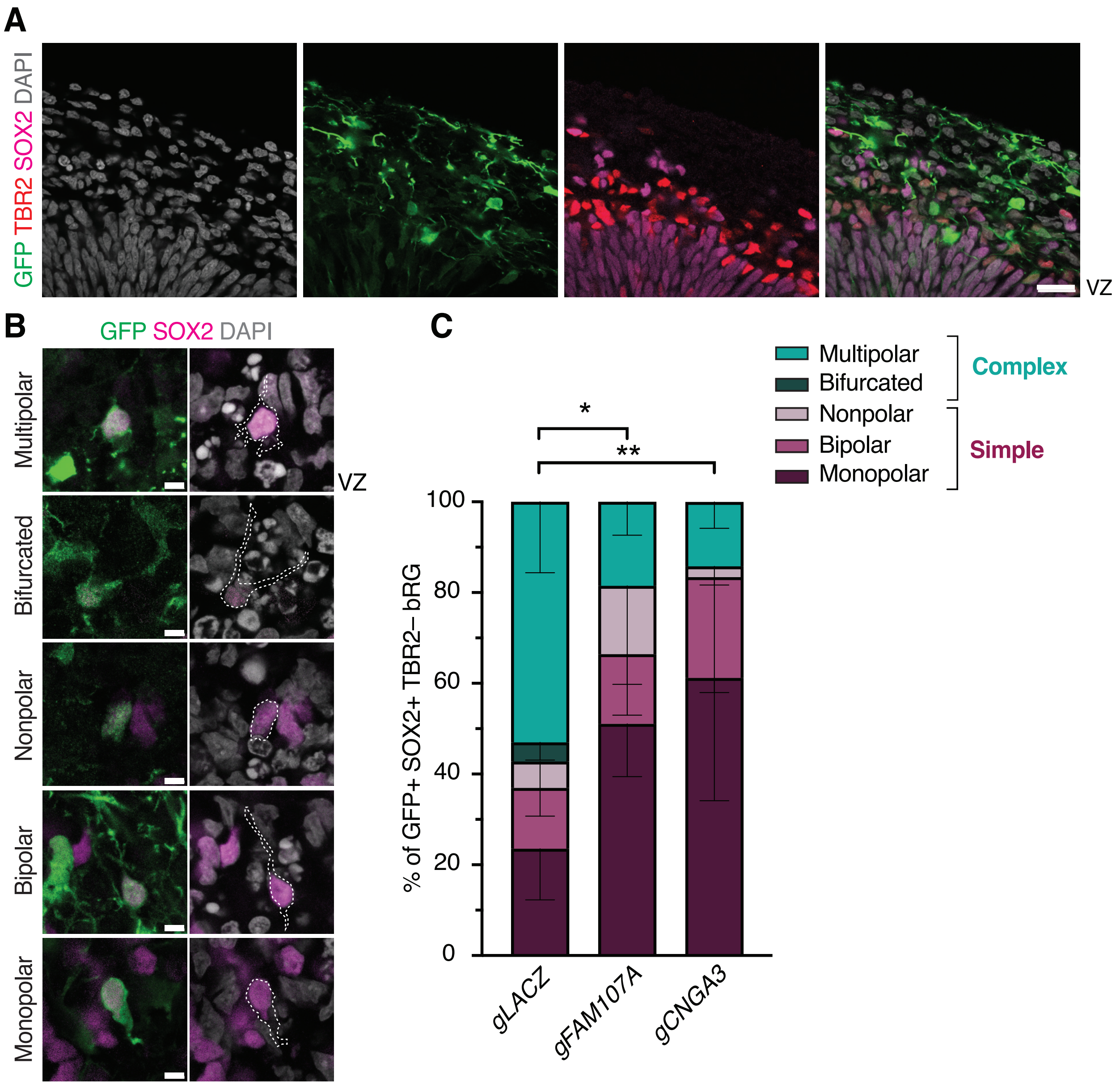


**Fig. S6.**

**Knockout of *FAM107A* and *CNGA3* leads to reduced morphological complexity of human bRG.**

(**A**) Overview image of electroporated hCOs showing immunofluorescence for GFP, SOX2, TBR2, and staining for DAPI. Basal radial glia identified as GFP+ SOX2+ TBR2– cells in the SVZ were scored in (C). Scale bar, 20 µm (**B**) Representative images of bRG morphotypes classified according to (*34*). Immunofluorescence for GFP and SOX2, and staining for DAPI. Scale bars, 5 µm. (**C**) Distribution of bRG morphotypes. Mean of 3 independent hCO differentiation and KO experiments. Total number of cells scored: g*LacZ* (Control), 41; g*FAM107A*, 28; g*CNGA3*, 36. Error bars represent SD; ** $p < 0.01$, * $p < 0.05$; 2-way ANOVA with Bonferroni post hoc test. Note the absence of bifurcated cells upon *FAM107A* and *CNGA3* knockout.

### Table S1. Oligonucleotides.

| Oligo | Species | Sequence | Reference |
|---|---|---|---|
| Gene expression analysis by qPCR | | | |
| GAPDH_F | human | 5'-AAGGTGAAGGTCGGAGTCAA | (*10*) |
| GAPDH_R | human | 5'-AATGAAGGGGTCATTGATGG | (*10*) |
| qSOX2_1F | human | 5'-CCAAGATGCACAACTCGGAG | This paper |
| qSOX2_1R | human | 5'-GCTTAGCCTCGTCGATGAAC | This paper |
| qEOMES_4F | human | 5'-GAGCCCTCAAAGACCCAGAC | (*10*) |
| qEOMES_4R | human | 5'-TTGCAAAGGGGTTATGATCAATC | (*10*) |
| qNGN2_1F | human | 5'-TGGTACACGATTGCAAACGG | (*10*) |
| qNGN2_1R | human | 5'-CTACGGGTCTTCTTGATGCG | (*10*) |
| qDCX_1F | human | 5'-TGTGTTTATTGCCTGTGGTCC | (*10*) |
| qDCX_1R | human | 5'-CTGTGGCTGATGGGTTTCC | (*10*) |
| qTUBB3_1F | human | 5'-AGCGTCTACTACAACGAGGC | (*10*) |
| qTUBB3_1R | human | 5'-AAGAGATGTCCAAAGGCCCC | (*10*) |
| qGorillaEOMES_1F | gorilla | 5'-GAGCCCTCAAAGACCCAGAG | This paper |
| RNA-seq | | | |
| dT-primer | N/A | C6-aminolinker-AAGCAGTGGTATCAACGCAGAGTCGAC TTTTTTTTTTTTTTTTTTTTTTTTTTTTVN, N = random base; V = any base beside thymidine | |
| TSO-primer | N/A | AAGCAGTGGTATCAACGCAGAGTACATrGrGrG, where rG stands for ribo-guanosine | |
| UP-primer | N/A | AAGCAGTGGTATCAACGCAGAGT | |
| Luciferase Assay constructs | | | |
| hCNGA3_wt | human | ATGGTACCAAACTCACAAACAAAGCAAGGGAAAAATGAAGCAACAAAAGCAGGGATTTATTGAAAACGAAAGTACCCTCCACAGGATGTGAGCTGGCTGAGCATAGGGGCTCCAGAGCCCAGTTACAGAATTTTCTGGTGTTTAAACACCCTCTAGAGGTTTCTCATTGGCCGCTTGATGTACACCCCATGCAAATGAAGTAGTGGCCTGCAGTCAGTCTGATTGCTAGCAT | This paper |
| hCNGA3_mut | human | ATGGTACCAAACTCACAAACAAAGCAAGGGAAAAATGAAGCAACAAAAGCAGGGATTTATTGAAAACGAAAGTACCCTCCACAGGGTGTGAGCTGGCTGAGCATAGGGGCTCCAGAGCCCAGTTACAGAATTTTCTGGTGTTTAAACACCCTCTAGAGGTTTCTCATTGGCCGCTTGATGTACACCCCATGCAAATGAAGTAGTGGCCTGCAGTCAGTCTGATTGCTAGCAT | This paper |
| cCNGA3 | chimpanzee | ATGGTACCAAACTCACAAACAAAGCAAGGGAAAAATGAGGCAACAAAAGCAGGGATTTATTGAAAACGA | This paper |

| | | | |
|---|---|---|---|
| | | AAGTACCCTCCACAGG**G**TGTGAG<br>CTGGCTGAGCATAGGGGCTCAAG<br>AGCCCAGTTACAGAATTTTCTGGT<br>GTTTAAACACCCTCTAGAGGTTTC<br>TCATTGGCCACTTGATGTACACCC<br>CATGCAAATGAAGTAGTGGCCTG<br>CAGTCAGTCTGATTGCTAGCAT | |
| hFAM107A_wt | human | ATGGTACCGGTGCTTCTCTGCTAC<br>TGGAAGAAAGTCGTGTTTCTTTTT<br>CCCTTCTCATTTTCAGAAAAGAAG<br>TGTGAAAGGATCAAAAGAAAGGA<br>TTTGCTTAGATGTTAGTGTGGCCT<br>GATTAGGTCGGAATTAGCCTTTTT<br>CTCTTTTCACTGAGAGAAGCAGAG<br>ACCATCTGCGGAAATTATTTAAAT<br>TCATGCGCCTTCCAGGGAAGCTGT<br>TTTATTTGCGCTAGCAT | This paper |
| hFAM107A_mut | human | ATGGTACCGGTGCTTCTCTGCTAT<br>TGGAAGAAAGTCGTGTTTCTTTTT<br>CCCTTCTCATTTTCAGAAAAGAAG<br>TGTGAAAGGATCAAAAGAAAGGA<br>TTTGCTTAGATGTTAGTGTGGCCT<br>GATTAGGTCGGAATTAGCCTTTTT<br>CTCTTTTCACTGAGAGAAGCAGAG<br>ACCATCTGCGGAAATTATTTAAAT<br>TCATGCGCCTTCCAGGGAAGCTGT<br>TTTATTTGCGCTAGCAT | This paper |
| cFAM107A | chimpanzee | ATGGTACCGGTTCTTCTCTGCTAT<br>TGGAAGAAAGTCGTGTTTCTTTTT<br>CCCTTCTCATTTTCAGAAAAGAAG<br>TGTGAAAGGATCAAAAGAAAGGA<br>TTTGCTTAGATGTTAGTGTGGCCT<br>GATTAGGTCGGAATTAGCCTTTTT<br>CTCTTTTCACTGAGAGAAGCAGAG<br>ACCATCTGCGGAAATTATTTAAAT<br>TCATGTGCCTTCCAGGGAGGCTGT<br>TTTATTTGCGCTAGCAT | This paper |
| Guide RNAs for CRSIPR/Cas9 knockout (underlined sequence is PAM) | | | |
| g*CNAG3* KO1 | human | 5'-<br>TCAGTGATACCAACAGGCTGTGG | (*38*) |
| g*CNGA3* KO2 | human | 5'-<br>GACGACCACGCAGTTCAAGCTGG | This paper |
| g*FAM107A* KO1 | human | 5'-<br>TCTTGATGAGCTGGTTCCGCCGG | (*38*) |
| g*FAM107A* KO2 | human | 5'-<br>GTGTCCTAGAGCACCGCCGGCGG | (*38*) |
| g*LacZ* | N/A | 5'-<br>TGCGAATACGCCCACGCGATCGG | (*39*) |
| PCR primers for gRNA validation | | | |
| CNGA3_Fwd | human | 5'- TGCATTTCCTGTAGTAATGG | This paper |
| CNGA3_Rev | human | 5'- GTCAAACCACCGGATAACC | This paper |
| FAM107A_Fwd | human | 5'- CTGGTTCAAATCACTCACCC | This paper |
| FAM107A_Rev | human | 5'- GGTGGGGAGAAATAGATGC | This paper |

### Table S2. Reagents and resources.

| **Reagent or Resource** | **Source** | **Identifier** |
|---|---|---|
| Antibodies | | |
| Rat anti-Ctip2 | Abcam ab18465 | RRID:AB_2064130 |
| Rabbit anti-Tbr2 | Abcam ab23345 | RRID:AB_778267 |
| Rabbit anti-Pax6 | Abcam ab195045 | RRID:AB_2750924 |
| Rabbit anti-Foxg1 | Abcam ab18259 | RRID:AB_732415 |
| Goat anti-Sox2 (1:200) | R&D Systems AF2018 | RRID:AB_355110 |
| Chicken anti-GFP (1:2,000) | Abcam ab13970 | RRID:AB_300798 |
| ZO-1 Monoclonal Antibody (ZO1-1A12), Alexa Fluor™ 647 | Invitrogen MA3-39100-A647 | RRID:AB_2663167 |
| FITC Anti-Ctip2 antibody [25B6] | Abcam ab123449 | RRID:AB_10973033 |
| V450 Mouse anti-Sox2 | BD Biosciences 561610 | RRID:AB_10712763 |
| PE Mouse anti-Sox2 | BD Biosciences 562195 | RRID:AB_10895118 |
| Alexa Fluor 488 Mouse anti-Sox2 | BD Biosciences 561593 | RRID:AB_10894382 |
| EOMES Monoclonal Antibody (WD1928), PE | Invitrogen 12-4877-42 | RRID:AB_2572615 |
| Donkey anti-Rat IgG, Alexa Fluor 488 conjugated | Invitrogen A-21208 | RRID:AB_141709 |
| Goat anti-Rabbit IgG, Alexa Fluor 405 conjugated | Invitrogen A-31556 | RRID:AB_221605 |
| Donkey anti-Goat IgG, Alexa Fluor 647 conjugated | Invitrogen A-21447 | RRID:AB_2535864 |
| Donkey anti-Rabbit IgG, Alexa Fluor 555 conjugated | Invitrogen A-31572 | RRID:AB_162543 |
| Donkey anti-Chicken IgG, Alexa Fluor 488 conjugated | Jackson Immuno Research 703-545-155 | RRID:AB_2340375 |
| Bacterial and virus strains | | |
| Subcloning Efficiency DH5α competent cells | Invitrogen | Cat. # 18265017 |
| Biological samples | | |
| GW13/14 human fetal brain tissue | This study | N/A |
| Chemicals, peptides, and recombinant proteins | | |
| DiI | Invitrogen | Cat. #D3911 |
| CellTracker CM-DiI Dye | Thermo Scientific | Cat. # C7000 |
| DAPI | Roche | Cat. # 10236276001 |
| Dimethyl Sulfoxide (DMSO) | Sigma Aldrich | Cat. # D2650-100ML |
| Tyrode's Salts (TS) | Sigma Aldrich | Cat. # T2145 |
| 36.5% formaldehyde solution | Sigma Aldrich | Cat. # F8775 |
| Glycine | Sigma Aldrich | Cat. # G8898 |
| Saponin | Sigma Aldrich | Cat. # 47036 |
| RNasin Ribonuclease Inhibitor | Promega | Cat. # N2165 |
| Bovine Serum Albumin (BSA) | Sigma Aldrich | Cat. # A2153 |
| EDTA disodium salt dihydrate | Carl Roth | Cat. # 8043.1 |
| Random Hexamers | Thermo Scientific | Cat. # N8080127 |
| 10 mM dNTPs | Thermo Scientific | Cat. # R0191 |
| Triton X-100 | Sigma Aldrich | Cat. # T9284 |
| RNaseOUT Recombinant Ribonuclease Inhibitor | Invitrogen | Cat. # 10777019 |
| SuperScript III Reverse Transcriptase, 5x First Strand Buffer, 100 mM DTT | Thermo Scientific | Cat. # 18080093 |
| LightCycler 480 SYBR Green I Master | Roche | Cat. # 06 991 076 702 |
| NP40 | Sigma/Roche | Cat. # 11332473001 |
| Tween-20 | Sigma/Roche | Cat. # 11332465001 |

| | | |
|---|---|---|
| Digitonin | Promega | Cat. # G944A |
| Nextera DNA Flex Library Prep Kit | Illumina | Cat. # FC-121-1030 |
| Proteinase K | Thermo Scientific | Cat. # EO0491 |
| NEBNext High- Fidelity 2x PCR Master Mix | New England Biolabs | Cat. # M0541S |
| SYBR Green I | Thermo Scientific | Cat. # S7563 |
| AMPure XP beads | Beckman Coulter | Cat. # A63880 |
| NovaSeq 6000 S4 Reagent Kit v1.5 (200 cycles) | Illumina | Cat. # 20028313 |
| SuperScript II Reverse Transcriptase, 5x First Strand Buffer, 100 mM DTT | Thermo Scientific | Cat. # 18064071 |
| KAPA HiFi HotStart ReadyMix | Roche | Cat. # 9420398001 |
| Illumina DNA Prep, (M) Tagmentation | Illumina | Cat. # 20060059 |
| RNase Inhibitor, Murine | New England Biolabs | Cat. # M0314L |
| Sera-Mag SpeedBeads | Cytvia | Cat. # GE24152105050250 |
| Heparin sodium salt from porcine intestinal mucosa | Sigma Aldrich | Cat. # H4784 |
| Trypsin-EDTA | Gibco | Cat. # 25300054 |
| Collagenase Type IV | Gibco | Cat. # 17104019 |
| DMEM/F12 | Gibco | Cat. # 31330 |
| Neurobasal | Gibco | Cat. # 21103049 |
| OptiMEM | Gibco | Cat. # 31985070 |
| Penicillin/ Streptomycin | Gibco | Cat. # 15140122 |
| Fetal Bovine Serum (FBS) | Sigma | Cat. #F7524 |
| MEM Non-Essential Amino Acids Solution (100X) | Gibco | Cat. # 11140050 |
| GlutaMAX Supplement | Gibco | Cat. # 35050038 |
| B-27 Supplement (50X), serum-free with VitA | Gibco | Cat. # 17504044 |
| B-27 Supplement (50X), minus VitA | Gibco | Cat. # 12587010 |
| N-2 Supplement (100X) | Gibco | Cat. # 17502048 |
| β-Mercaptoethanol | Gibco | Cat. # 21985023 |
| TrypLE Express Enzyme (1x), no phenol red | Gibco | Cat. # 12604021 |
| ReLeSR | Stem Cell Technologies | Cat. # 05873 |
| Rock inhibitor, Y-27632 (2HCl) | Stem Cell Technologies | Cat. # 72308 |
| mTeSR1 Complete Kit | Stem Cell Technologies | Cat. # 85850 |
| StemFlex Medium | Gibco | Cat. # A3349401 |
| Matrigel hESC-Qualified Matrix, LDEV-free | Corning | Cat. # 354277 |
| STEMdiff Cerebral Organoid Kit | Stem Cell Technologies | Cat. # 8570 |
| STEMdiff Cerebral Organoid Maturation Kit | Stem Cell Technologies | Cat. # 8571 |
| Amphotericin B (Fungizone) | Gibco | Cat. # 11520496 |
| KnockOut Serum Replacement | Gibco | Cat. # 10828010 |
| Insulin Solution, Human recombinant | Sigma Aldrich | Cat. # I9278 |
| A83-01 | Stem Cell Technologies | Cat. # 72022 |
| Dorsomorphine | Stem Cell Technologies | Cat. # 72102 |
| CHIR-99021 | Stem Cell Technologies | Cat. # 72052 |
| SB-431542 | Stem Cell Technologies | Cat. # 72232 |
| Recombinant Human/Murine/Rat BDNF | Gibco | Cat. # 450-02 |
| Recombinant Human GDNF | Gibco | Cat. # 450-10 |
| Ascorbic acid | Sigma Aldrich | Cat. # 1043003 |
| Dibutyryl-cAMP | Stem Cell Technologies | Cat. # 73882 |
| UltraPure low melting point agarose | Invitrogen | Cat. # 16520100 |

| | | |
|---|---|---|
| Matrigel Growth Factor Reduced (GFR) Basement Membrane Matrix, LDEV-free | Corning | Cat. # 354230 |
| Trypan Blue Solution, 0.4% | Gibco | Cat. # 15250061 |
| SPRIselect Beads | Beckman Coulter | Cat. # B23317 |
| KpnI-HF | New England Biolabs | Cat. # R3142S |
| BmtI-HF | New England Biolabs | Cat. # R3658S |
| T4 DNA Ligase | New England Biolabs | Cat. # M0202 |
| Quick CIP Phosphatase | New England Biolabs | Cat. # M0525 |
| Horse Serum, heat inactivated | Gibco | Cat. # 26050088 |
| Mowiol 4-88 | Sigma Aldrich | Cat. # 81381 |
| Critical commercial assays | | |
| MACS Neural Tissue Dissociation Kit (P) | Miltenyi Biotec | Cat. # 130-092-628 |
| Quick-RNA FFPE MiniPrep Kit | Zymo Research | Cat. # R1008 |
| DNA Clean and Concentrator-5 Kit | Zymo Research | Cat. # D4014 |
| Lipofectamine 2000 | Invitrogen | Cat. # 11668027 |
| QIAprep Spin Miniprep Kit | Qiagen | Cat. # 27106 |
| Dual-Luciferase Reporter Assay System | Promega | Cat. # E1910 |
| P3 Primary Cell 4DNucleofector™ X Kit S | Lonza | Cat. # V4XP-3032 |
| QIAquick Gel Extraction Kit | Qiagen | Cat. # 28706 |
| Experimental models: Cell lines | | |
| Human: UCSFi001-A / WTC11 | (*1*) | RRID:CVCL_Y803 |
| Human: CRTDi004-A | (*2*) | RRID:CVCL_YR23 |
| Gorilla: GC1 iPSC | (*4*) | N/A |
| Chimpanzee: SandraA iPSC | (*3*) | N/A |
| Mouse: Neuro-2A | | N/A |
| Oligonucleotides | | |
| Oligonucleotides, see Table S1 | This paper | N/A |
| Software and algorithms | | |
| Fiji/ImageJ | Fiji/ImageJ | https://imagej.nih.gov/ij/ |
| ZEN black (11.0.1.190) or ZEN blue (v2012) | Zeiss | N/A |
| Prism (8.4.3) | GraphPad software | N/A |
| Geneious Prime (2019.2.1) | Biomatters Ltd. | N/A |
| Adobe Illustrator (27.2) | Adobe | N/A |
| Affinity Photo + Designer (1.10.5.1342) | Serif Ltd. | N/A |
| FACSDiva (8.0.2 and 9.0.1) | BD Biosciences | N/A |
| FastQC (v0.11.6) | Babraham Bioinformatics | https://www.bioinformatics.babraham.ac.uk/projects/fastqc/ |
| STAR | (*29*) | |
| ENCODE ATAC-seq pipeline | (*18*) | |
| kallisto | (*32*) | |
| HALPER | (*19*) | |
| DESeq2 | (*22*) | |
| MonaLisa | (*28*) | |
| Other | | |
| Phylogenetic pictures | N/A | https://www.phylopic.org/ |
| Light-Cycler 480 Multiwell Plate 96, white | Roche | Cat. # 04729692001 |

| | | |
|---|---|---|
| DNA LoBind microcentrifuge tubes, 1.5 mL | Eppendorf | Cat. # 0030108051 |
| Falcon 5 mL Round Bottom Polystyrene Test Tube, with Snap Cap, Sterile | Corning | Cat. # 352058 |
| Falcon 5 mL Round Bottom Polystyrene Test Tube, with Cell Strainer Snap Cap | Corning | Cat. # 352235 |
| 96-well, white wall, clear bottom plates | Greiner | Cat. # 655095 |
| Nunc Cell-Culture Treated Multidishes, 6-well | Thermo Scientific | Cat. # 140675 |
| 96-well Clear Round Bottom Ultra-Low Attachment Microplate, with lid, sterile | Corning | Cat. # 7007 |
| Nunc IVF Dishes, 54 mm diameter | Thermo Scientific | Cat. # 150270 |
| 6-well Clear Flat Bottom Ultra-Low Attachment Well Plates, Sterile | Corning | Cat. # 3471 |
| pGL4.23[luc2/minP] | Promega | GenBank: DQ904455.1 |
| pRL-CMV | Promega | GenBank: AF025843.2 |
| pCAG-GFP | (*40*) | Addgene #11160 |

27. C. Gene Ontology, S. A. Aleksander, J. Balhoff, S. Carbon, J. M. Cherry, H. J. Drabkin, D. Ebert, M. Feuermann, P. Gaudet, N. L. Harris, D. P. Hill, R. Lee, H. Mi, S. Moxon, C. J. Mungall, A. Muruganugan, T. Mushayahama, P. W. Sternberg, P. D. Thomas, K. Van Auken, J. Ramsey, D. A. Siegele, R. L. Chisholm, P. Fey, M. C. Aspromonte, M. V. Nugnes, F. Quaglia, S. Tosatto, M. Giglio, S. Nadendla, G. Antonazzo, H. Attrill, G. Dos Santos, S. Marygold, V. Strelets, C. J. Tabone, J. Thurmond, P. Zhou, S. H. Ahmed, P. Asanitthong, D. Luna Buitrago, M. N. Erdol, M. C. Gage, M. Ali Kadhum, K. Y. C. Li, M. Long, A. Michalak, A. Pesala, A. Pritazahra, S. C. C. Saverimuttu, R. Su, K. E. Thurlow, R. C. Lovering, C. Logie, S. Oliferenko, J. Blake, K. Christie, L. Corbani, M. E. Dolan, H. J. Drabkin, D. P. Hill, L. Ni, D. Sitnikov, C. Smith, A. Cuzick, J. Seager, L. Cooper, J. Elser, P. Jaiswal, P. Gupta, P. Jaiswal, S. Naithani, M. Lera-Ramirez, K. Rutherford, V. Wood, J. L. De Pons, M. R. Dwinell, G. T. Hayman, M. L. Kaldunski, A. E. Kwitek, S. J. F. Laulederkind, M. A. Tutaj, M. Vedi, S. J. Wang, P. D'Eustachio, L. Aimo, K. Axelsen, A. Bridge, N. Hyka-Nouspikel, A. Morgat, S. A. Aleksander, J. M. Cherry, S. R. Engel, K. Karra, S. R. Miyasato, R. S. Nash, M. S. Skrzypek, S. Weng, E. D. Wong, E. Bakker, T. Z. Berardini, L. Reiser, A. Auchincloss, K. Axelsen, G. Argoud-Puy, M. C. Blatter, E. Boutet, L. Breuza, A. Bridge, C. Casals-Casas, E. Coudert, A. Estreicher, M. Livia Famiglietti, M. Feuermann, A. Gos, N. Gruaz-Gumowski, C. Hulo, N. Hyka-Nouspikel, F. Jungo, P. Le Mercier, D. Lieberherr, P. Masson, A. Morgat, I. Pedruzzi, L. Pourcel, S. Poux, C. Rivoire, S. Sundaram, A. Bateman, E. Bowler-Barnett, A. J. H. Bye, P. Denny, A. Ignatchenko, R. Ishtiaq, A. Lock, Y. Lussi, M. Magrane, M. J. Martin, S. Orchard, P. Raposo, E. Speretta, N. Tyagi, K. Warner, R. Zaru, A. D. Diehl, R. Lee, J. Chan, S. Diamantakis, D. Raciti, M. Zarowiecki, M. Fisher, C. James-Zorn, V. Ponferrada, A. Zorn, S. Ramachandran, L. Ruzicka, M. Westerfield, The Gene Ontology knowledgebase in 2023. *Genetics* **224**, (2023).
